# Individual differences in probabilistic learning and updating predictive representations in individuals with obsessive-compulsive tendencies

**DOI:** 10.1101/2024.12.03.626598

**Authors:** Bianka Brezóczki, Bence Csaba Farkas, Flóra Hann, Orsolya Pesthy, Eszter Tóth-Fáber, Kinga Farkas, Katalin Csigó, Dezső Németh, Teodóra Vékony

## Abstract

Obsessive-compulsive (OC) tendencies involve intrusive thoughts and rigid, repetitive behaviours that also manifest at the subclinical level in the general population. The neurocognitive factors driving the development and persistence of the excessive presence of these tendencies remain highly elusive, though emerging theories emphasize the role of implicit information processing. Despite various empirical studies on distinct neurocognitive processes, the incidental retrieval of environmental structures in dynamic and noisy environments, such as probabilistic learning, has received relatively little attention. In this study, we aimed to unravel potential individual differences in implicit probabilistic learning and the updating of predictive representations related to OC tendencies in the general population. We conducted two independent online experiments (N_Study1_ = 164, N_Study2_ = 257) with young adults. Probabilistic learning was assessed using a reliable implicit visuomotor probabilistic learning task, which involved sequences with second-order non-adjacent dependencies. Our findings revealed that even among individuals displaying a broad spectrum of OC tendencies within a non-clinical population, implicit probabilistic learning remained remarkably robust. Furthermore, the results highlighted effective updating capabilities of predictive representations, which were not influenced by OC tendencies. These results offer new insights into individual differences in probabilistic learning and updating in relation to OC tendencies, contributing to theoretical, methodological, and practical approaches for understanding the maladaptive behavioural manifestations of OC disorder and subclinical tendencies.

## Introduction

During their morning routines, many people repeatedly check whether they have locked the front door or confirm that their wallet and phone are in their bag or pockets. Over time, these may become automatic, habitual actions. Such behaviours often reflect non-clinical tendencies of Obsessive-Compulsive Disorder (OCD) even in healthy individuals (1). OCD is defined by the excessive and impeding presence of intrusive thoughts (‘obsessions’) and rigid, repetitive actions (‘compulsions’) (2). Neurobiological models underline structural and functional alterations in the frontostriatal circuits as key biomarkers of the disorder (3–11). These brain regions are engaged in creating stimulus-response associations (3,12), which are sustained and reinforced over time, influencing the development and persistence of compulsive behaviours (13,14). This feedback-sensitive nature of OCD led previous research to focus primarily on supervised learning processes, where learning is reinforced with feedback or reward (1,15–19). However, associative contingencies can be incidentally learned from environmental cues, even without supervision, serving as a key component of predictive processes (20–22). This type of unsupervised learning remains poorly understood in the context of OCD as well as in subclinical obsessive-compulsive (OC) tendencies. Therefore, the study aims to address this gap.

In supervised learning or reinforcement learning, participants receive feedback or reward for their performance throughout the task. Such paradigms used in OCD research are the probabilistic classification task, where participants make predictions based on cues that are probabilistically linked to outcomes (17,18,23–25) and associative supervised learning tasks mainly reflect the learning of deterministic cue-outcome contingencies (13,14). Previous studies demonstrated intact perfomance on both probabilistic suprvised learning and associative supervised learning tasks in OCD (17,18,23, for exceptions see: 24,25). However unsupervised learning paradigms showed weaker performance (26–32, for exceptions see: 33– 36). Most of these unsupervised studies focused on visuomotor sequence learning, but the application of fixed deterministic contingencies has recently been emphasized as a key weakness of these studies (for a review, see: (26). As we discern, the existing paradigms vary in the presence or absence of supervision, as well as in the predictability of the stimulus-response contingencies. However, one indispensable aspect has been overlooked so far. Namely, that our brain can implicitly extract hidden probabilistic patterns embedded in noisy and less relevant stimuli without supervision, allowing us to form predictive representations (22). Therefore, in this study, we aim to overcome previous limitations by focusing on probabilistic contingencies rather than deterministic ones among OC tendencies – as proposed earlier (26) – in an unsupervised manner.

Since our environment is constantly changing, in addition to the extraction of hidden probabilistic contingencies, it is also essential to assess the stability of the established predictive representations and our ability to effectively update them in response to volatile situations. Prominent theories based on supervised methods propose that individuals with OCD tend to rigidly stick to previously learned information and find it difficult to update their behaviour when facing a novel condition (13,14,37). Another recent theory similarly emphasizes the role of uncertainty and pervasive doubt about previously acquired knowledge, leading individuals to rely on the most recent outcomes, thereby generating highly exploratory and distrustful behaviour (38,39). This uncertainty can encompass not just the clinical OCD population, but subclinical OC tendencies (39). Both theories imply that individuals with OCD or OC tendencies face challenges in updating their behaviour, however, neither theoretical model elucidates the updating of predictive representations arising from the incidental learning of probabilistic contingencies. Actually, at times, the environment may change in subtle, unconscious ways without any prior warning, prompting individuals to update their behaviour without realizing it. This requires the ability to adjust predictive representations accordingly. Nonetheless, this ability has not yet been examined in the context of OCD and OC tendencies. Thus, our question is: to what extent do OC tendencies contribute to the resistance and updating capability of predictive representations in a volatile environment, in an unsupervised manner?

Investigating learning and updating in OCD is crucial, and most of the aforementioned findings have been conducted primarily with clinically diagnosed OCD patients. However, numerous studies increasingly support a dimensional understanding of OCD over dichotomous categories (1,15,40–43). This approach emphasizes the presence of subclinical OC tendencies or traits in the general population and their manifestations in behavioural variability and brain structures (1,15,40–43). Furthermore, it has been suggested that examining non-clinical samples could offer insights into aspects of the continuum that connect subclinical OC tendencies to clinically severe OC symptoms (40). In this regard, we lack information on how OC tendencies are associated with potential individual differences in the acquisition of probabilistic contingencies and the resilience or updating of predictive representations. Moreover, to overcome the limitations of prior studies, we aim to assess implicit probabilistic processes using an ecologically valid visuomotor statistical learning task with second-order non-adjacent dependencies, which examines learning of probabilistic contingencies in the absence of supervision regarding learning performance (44–47). We conducted two online experiments involving relatively large, general population samples (N_Study1_ = 164, N_Study2_ = 257). In both studies, OC tendencies are handled as a continuous spectrum, ranging from the lowest to the highest prevalence of OC tendencies. In Study 1, we aimed to pinpoint how OC tendencies contribute to potential individual differences in probabilistic learning, while in Study 2, to the resistance and updating of predictive representations in adjusting to a volatile condition.

## Methods

### Participants

Study 1 involved 224 university students who participated in an online experiment, for which they received course credit. To ensure data integrity, stringent quality control measures were employed, resulting in the exclusion of 60 participants who did not comply with instructions. We aimed to meet multiple criteria for data quality control, which is detailed in the flow chart of participant exclusion after data collection as illustrated in Supplementary Materials S1. This led us to conduct our analysis on 73.21% of the initial participants in Study 1. The final sample of Study 1 contained 164 participants (111 female, 48 male, and 2 non-specified, 3 declined to answer; M_Age_=21.76 years; SD_Age_=5.21).

In Study 2, 344 university students participated in exchange for course credit. Similar to Study 1, we ensured data integrity (see Supplementary Materials S1 for more details), leading to the exclusion of 87 participants, thus yielding a final sample of 257 (74.7%) individuals (189 female, 65 male, and 1 non-specified, M_Age_=22.47 years; SD_Age_=5.39). Participants in both studies gave informed consent, and the studies obtained approval from the Research Ethics Committee of Eötvös Loránd University Budapest, Hungary (2021/504), under the principles of the Declaration of Helsinki.

### Measures

#### Alternating Serial Reaction Time task

To assess probabilistic learning, we employed a visuomotor learning paradigm, the Alternating Serial Reaction Time (ASRT) task (44,46). The task was created in JavaScript using the jsPsych library version 6.1.0 (48,49). Participants encountered a visual stimulus, represented by a drawing of a dog’s head, displayed in one of four positions arranged horizontally on the computer screen. Their goal was to quickly and accurately identify the location of the target stimulus by pressing the corresponding key on the keyboard (“S”, “F”, “J”, or “L” keys from left to right, respectively) (Fig. 1A). Stimuli remained visible until a correct response was provided, followed by a 120 ms response to stimulus interval before the next stimulus appeared. In the event of an incorrect response, the target stimulus remained present until a correct response was provided. Without the participants’ awareness, the stimuli adhered to a probabilistic eight-element sequence, alternating between pattern and random elements (e.g., 2 – r – 4 – r – 3 – r – 1 – r, where “r” denotes a random location, and the numbers signify predetermined positions from left to right). The task incorporated an alternating structure where some runs of three consecutive elements were more predictable (high-probability triplets) than others (low-probability triplets). A trial denotes a single element within the sequence, which can be either a pattern or a random element, and notably, it serves as the final element in a high- or low-probability triplet. For instance, in a sequence like 2–r–4–r–3–r–1–r, triplets such as 2-X-4, 4-X-3, 3-X-1, and 1-X-2 (where X represents the middle element of a triplet) occurred more frequently than triplets such as 2-X-1 or 3-X-2, as their first and third elements could belong to either a pattern or random trial (Fig. 1B). High-probability triplets could be produced through the inclusion of either two pattern trials and one random trial in the center (occurring in 50% of trials), or by incorporating two random trials and one pattern trial in the center (occurring in 12.5% of trials). The remaining trials were categorized as low-probability triplets, with the final element appearing as random (occurring in 37.5% of trials) (Fig. 1C).

**Figure 1.**
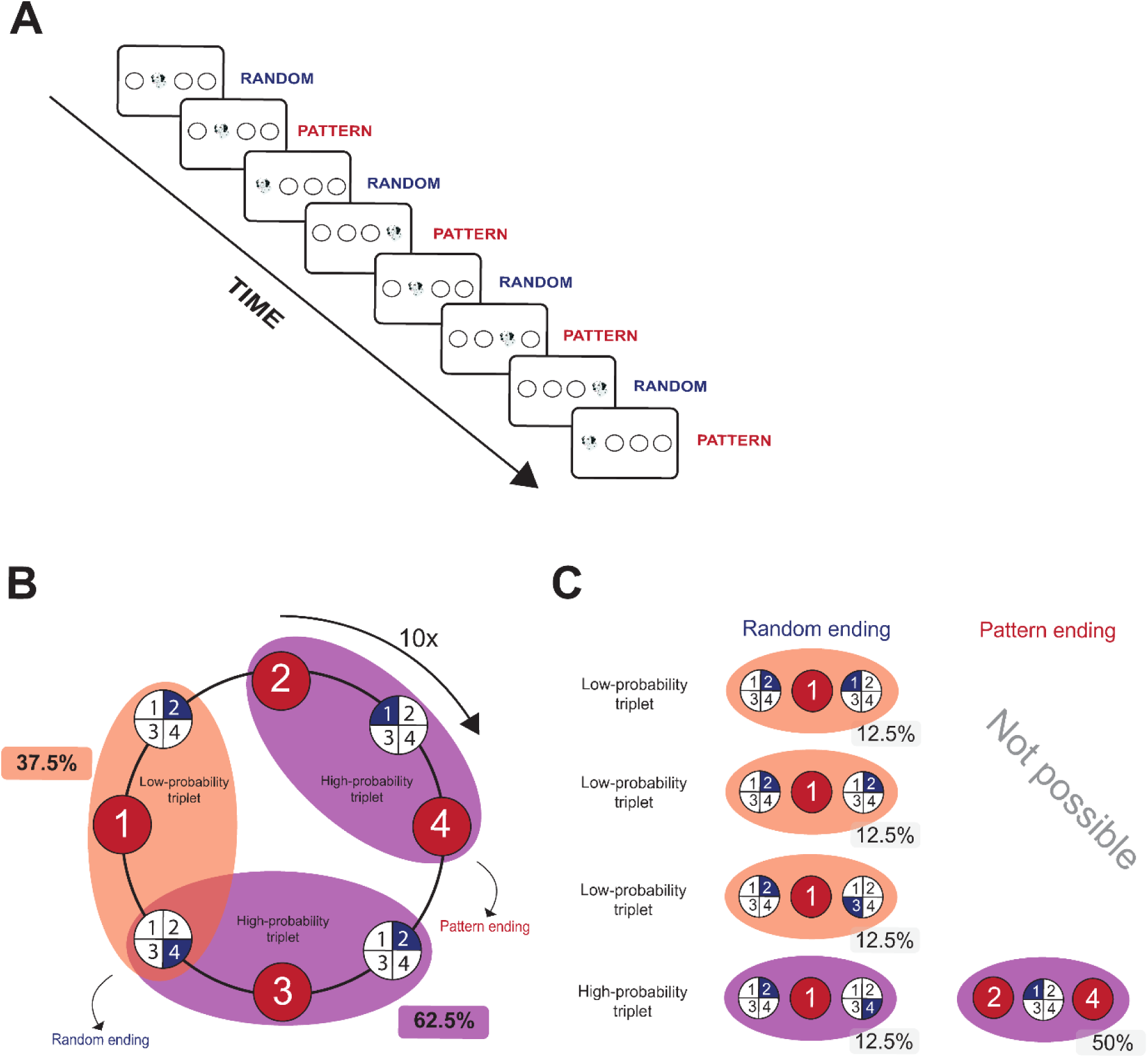
ASRT Task Structure. **A.)** In the ASRT task, participants were required to press keys that matched the location of a target stimulus, represented by a dog’s head. Every trial followed an 8-element probabilistic sequence. This sequence included random elements interspersed among the patterned elements, forming a series like 2-r-4-r-3-r-1-r, where the numbers indicate the locations of the pattern trials from left to right, and ‘r’ signifies random positions. **B.)** Formation of triplets in the ASRT task. Pattern elements, which consistently appear in the same positions throughout the task, are represented with red backgrounds. Random elements, chosen randomly from four possible positions, are shown with blue backgrounds. Each trial was categorized as the third element in a sequence of three consecutive trials (a triplet). The probabilistic sequence structure resulted in some triplets occurring more frequently (high-probability triplets) than others (low-probability triplets). For visualization purposes, only three triplets are highlighted in this subfigure: 2(P)-1(R)-4(P) as a pattern-ending high-probability triplet, 2(R)-3(P)-4(R) as a random-ending high-probability triplet, and 4(R)-1(P)-2(R) as a random-ending low-probability triplet. **C.)** The formation of high-probability triplets could involve the occurrence of either two pattern trials and one random trial at the center, which occurred in 50% of trials, or two random trials and one pattern trial at the center, which occurred in 12.5% of trials. In total, 62.5% of all trials constituted the final element of a high-probability triplet, while the remaining 37.5% were the final elements of a low-probability triplet.

In Study 1, the ASRT task comprised 25 blocks, encompassing 10 repetitions of an eight-element sequence, resulting in 80 stimuli within each block. After completing each block, participants received individual feedback on their average accuracy and reaction time, but no feedback was given on their learning performance (as they were not aware that the task involved the learning of a regularity).

In Study 2, we utilized the same task structure but modified the task design. In the first part, 15 blocks were recorded, resulting in 1200 trials (hereafter referred to as the Learning Phase). During this phase, participants learned and practiced a predetermined probabilistic sequence (Sequence A: e.g., 2-r-4-r-3-r-1-r). This phase was followed by a delay period (M = 46.08 min, SD = 19.29). After the break, the second part of the task commenced, involving another 15 blocks, from Block 16 to Block 30 (1200 trials). In Block 16-20, participants were reintroduced to the original, well-learned sequence from the previous phase (Sequence A). However, in block 21-25, without any prior warning, we changed the sequence structure (i.e., Interference phase). We gave participants the reversed version of the original sequence (e.g., Sequence A: 2-r-4-r-3-r-1-r, Sequence B: 1-r-3-r-4-r-2-r). In block 26-30, participants were presented the original sequence (A) again (Fig. 2B). The structural change in block 21-25 resulted in some alterations in the probability of the triplets. Triplets that were high-probability during the learning phase became low-probability during the interference phase (‘H L’), or they were low-probability during the learning phase that became high-probability during the Interference phase (‘L H’). There were also triplets that were low-probability in both conditions (‘L L’). Due to the reversed nature of the sequence and the lack of overlap, the formation of ‘H H’ triplets was not possible (47). With this design, in addition to assessing probabilistic learning and visuomotor performance improvement, we can also measure sensitivity to interfering information. Based on Horváth et al. (2022), this can be achieved in two ways: the difference between performance on ‘H L’ and ‘L L’ triplets reflects the resistance of previously well-learned information to new, interfering information (referred to as ‘Old knowledge’). Whereas the difference between performance on ‘L H’ and ‘L L’ triplets indicates the robustness of new information compared to previously learned information, so-called ‘New knowledge’ (Fig. 2C).

**Figure 2.**
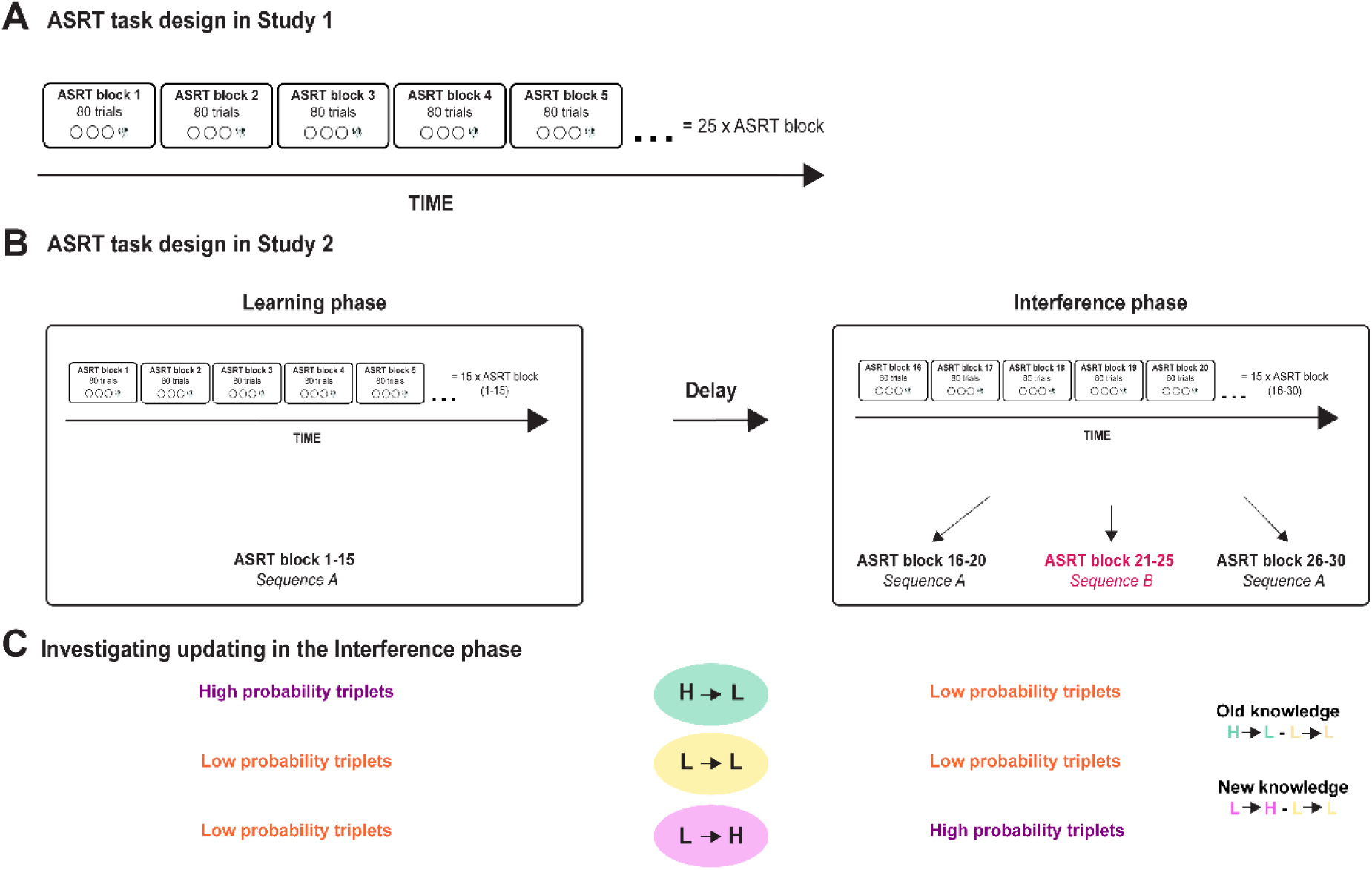
ASRT Task Experimental Design. **A.)** ASRT task design in Study 1. In Study 1, the ASRT task comprised 25 blocks, encompassing 10 repetitions of an eight-element sequence, resulting in 80 stimuli within each block. **B.)** The ASRT task design in Study 2 involved multiple phases. In the Learning phase, participants were given the same sequence (Sequence A) across 15 blocks, with each block containing 80 trials. This was followed by a delay (M = 46.08 min, SD = 19.29), during which participants completed questionnaires. In the Interference phase, participants first practiced Sequence A again in block 16-20. In block 21-25, the underlying sequence was changed without warning to the reversed version of the original sequence (Sequence B). Finally, in block 26-30, participants were re-exposed to Sequence A. **C.)** Investigating updating in the Interference phase in Study 2. The structural change in block 21-25 resulted in changes in the probability of certain triplets. Triplets that were high-probability during the Learning phase became low-probability during the Interference phase (‘H L’), while those that were low-probability during the Learning phase became high-probability during the Interference phase (‘L H’). Additionally, some triplets remained low-probability in both phases (‘L L’). The difference between the frequencies of ‘H L’ and ‘L L’ triplets reflects the resistance of previously well-learned probabilistic information to new interfering information, referred to as ‘Old knowledge’. Conversely, the difference in probabilities between ‘L H’ and ‘L L’ triplets indicates the robustness of new probabilistic information compared to previously learned information, referred to as ‘New knowledge’.

**Figure 3.**
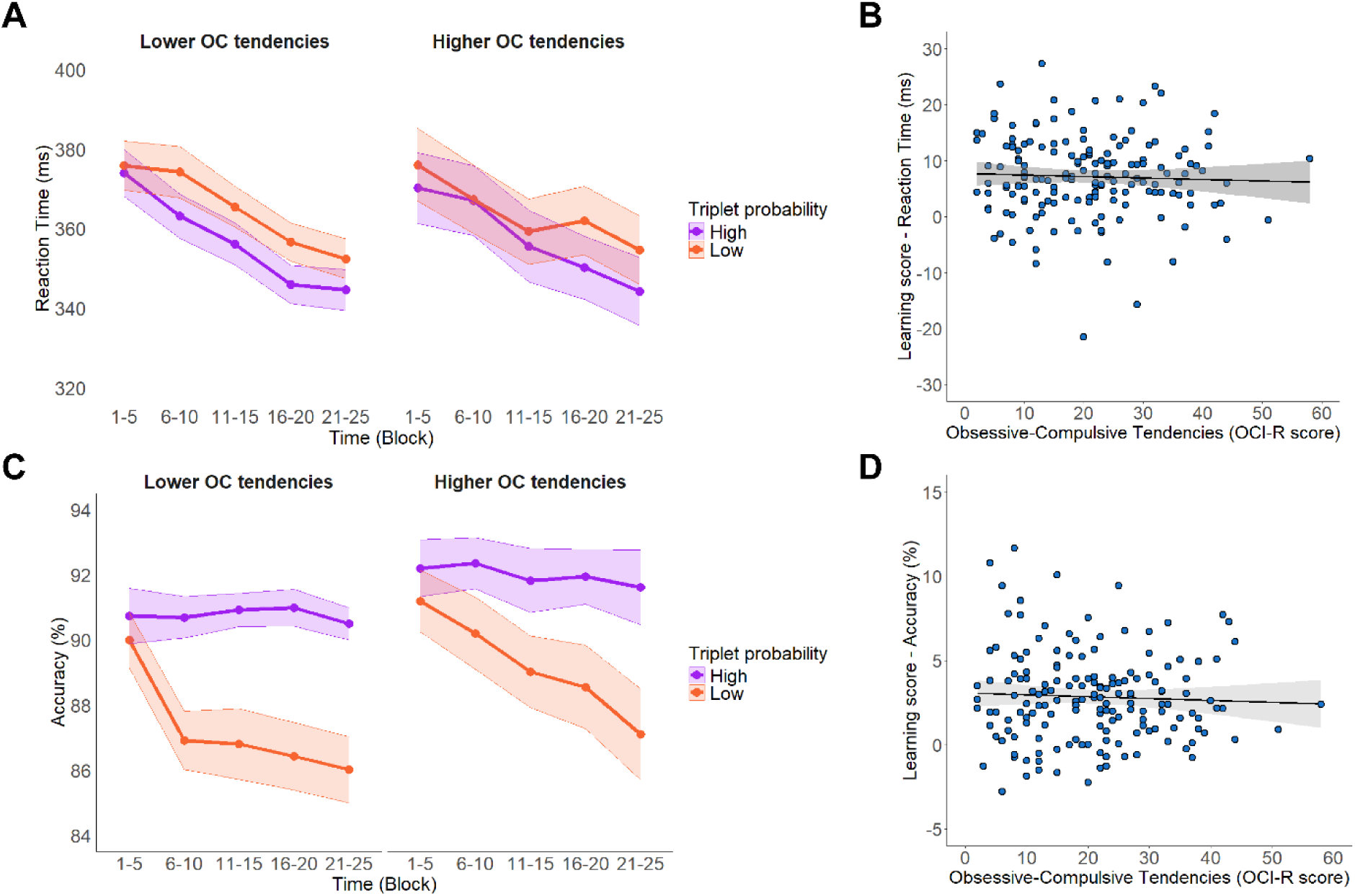
Probabilistic learning performance metrics as a function of OC tendencies in Study 1. **A.** Probabilistic learning performance in terms of temporal trajectory (blockwise) reaction time. The orange line represents the responses given to the low-probability triplets and the purple line represents responses given to the high-probability triplets. Note that OC tendencies measured by OCI-R score was treated as a continuous variable in our analyses, but for more illustrative data visualization, we depicted results at Lower OC tendencies (OCI-R score: 20.73-11.33, on the left) and Higher OC tendencies (OCI-R score: 20.73 +11.33, on the right) based on standard deviations from the mean. The x-axis indicates Time (Block 1-5, Block 6-10, Block 11-15, Block 16-20, Block 21-25), y-axis indicates RT (in ms). Participants also showed evidence of implicit SL, as they increasingly became faster for high-probability triplets than for low-probability triplets as the task progressed. However, OC tendencies did not influence either the amount, or the trajectory of learning. In all subfigures, points indicate means and error bars indicate SEM. **B.** Correlation between OC tendencies and difference learning scores based on reaction time for low- and high-probability triplets for the entire task. The correlation between probabilistic learning and OC tendencies in reaction time was not significant (*r*(162) = −.029, *p* = .709), indicating that OC tendencies do not relate to probabilistic learning as measured by reaction time. **C**. Probabilistic learning performance in terms of temporal trajectory (blockwise) mean accuracy. The orange line represents the responses given to the Low probability triplets and the purple line represents responses given to the high-probability triplets. Note that OC tendency was treated as a continuous variable in our analyses, but for more illustrative data visualization, we demonstrated results at Lower OC tendencies (OCI-R score: 20.73-11.33, on the left) and Higher OC tendencies (OCI-R score: 20.73 +11.33, on the right) based on standard deviations from the mean. X-axis indicates Time (1-5), Y-axis indicates mean accuracy (in %). Participants also showed evidence of implicit probabilistic learning, as they increasingly became more accurate for high-probability triplets than for low-probability triplets as the task progressed. However, OC tendencies did not influence the amount, nor the trajectory of learning. In all subfigures, points indicate means and error bars indicate SEM. **D.** Correlation between OC tendencies and difference learning scores based on accuracy for high- and low-probability triplets for the entire task. The correlation between probabilistic learning and OC tendencies in accuracy was not significant (*r* (162) = −.035; *p* = .653), indicating that OC tendencies do not relate to probabilistic learning as measured by accuracy.

#### Obsessive-Compulsive Inventory-Revised

The evaluation of OC tendencies involved the use of a self-report questionnaire designed to assess both the severity and nature of these symptoms. This questionnaire, known as the Revised version of the Obsessive-Compulsive Inventory (OCI-R; (50), consists of 18 items rated on a 5-point Likert scale (0 = Not at all; 4 = Extremely). Participants were instructed to indicate how much each described experience bothered them over the preceding month. The questionnaire has shown robust psychometric characteristics in both clinical (50,51) and subclinical populations (52). The questionnaire has a maximum achievable score of 72 points (50–52). In Study 1, the highest score was 58 (M = 20.72, SD = 11.30), while in Study 2, it was 51 (M = 14.24, SD = 11.63).

#### Procedure

Both experiments were hosted by the Gorilla Experiment Builder platform (https://www.gorilla.sc) (53) The experiments began with participants reading an informational briefing and giving their consent to participation. Following this, Study 1 proceeded with the ASRT task, where participants completed two practice blocks before engaging in 25 subsequent task blocks. The initial practice blocks consisted entirely of random trials, which were not analyzed here. To evaluate participants’ perception of any discernible patterns during the task, we administered inquiries after its completion, prompting them to report any patterns they noticed throughout the task. Following this, participants were questioned regarding their awareness of the concealed sequence upon completion of the ASRT task. Specifically, they were asked whether they detected any regularities or patterns during the task and to provide details if they did. None of the participants could accurately describe the alternating sequence, which means that the task could remain entirely implicit, as confirmed earlier (47,54). Subsequently, we assessed the prevalence of OC tendencies with the OCI-R, as well as some demographic questions about socioeconomic status, sleep, health, and handedness. Considering the online nature of the experiment, participants were presented with additional brief questions regarding the circumstances of task completion. This was aimed at controlling for any potential confounding variables in our analyses (as seen in Supplementary Materials S1).

In Study 2, after obtaining informed consent, participants proceeded with the ASRT task, starting with the Learning phase. During this phase, participants began with two practice blocks to familiarize themselves with the task, followed by 15 blocks. The practice phase consisted of two blocks of random stimuli, which were not included in the analyses. A delay period followed the Learning phase, during which participants completed various questionnaires, including the OCI-R and demographic questions about socioeconomic status, sleep, health, and handedness — this period lasted 46.08 minutes on average (± 19.29 SD). Following this, participants entered the Interference Phase, where they did not perform any practice blocks and immediately proceeded to testing. After completing Block 30, participants were asked if they noticed any patterns during the task and, if so, were prompted to recall them. Similar to Study 1, participants were presented with the same additional brief questions about the conditions under which they completed the experiment, as detailed in the Supplementary Materials S1, to ensure the integrity of the experimental data.

#### Data manipulation and analysis

Data preparation, statistical analyses, and visualization were conducted using R 4.3.3. (55). Concerning the data of the ASRT task, first, we categorized each trial based on whether they were the last element of a high- or low-probability triplet. We excluded responses that were not the first one for the given trial. Trials below 150 ms and above 1000 ms were also removed, as well as the responses below and above three absolute median deviations. Repetitions (e.g., 2-2-2, 1-1-1) and Trills (e.g., 2-1-2, 3-2-3) were removed from the analysis since participants may show faster motor response inclinations on these types of trials (Soetens et al., 2004).

Furthermore, we removed the first two trials of each block, as they do not belong to any identified triplet type. Finally, incorrect responses were excluded from the reaction time (RT) analysis. In Study 1, these criteria necessitated the removal of 17.9% of trials for the learning analyses of accuracy and 25.5% for analyses of RTs. In the Learning phase of Study 2, 19.2% of trials for analyses of accuracy and 26.1% for analyses of RTs; and 16.7% of trials from the Interference phase for analyses of accuracy and 23.7% for analyses of RTs were removed.

##### Learning phase

We report performance indicators calculated from both RT and accuracy data. We introduce two distinct forms of learning: probabilistic learning and improvement in visuomotor performance. Probabilistic learning was calculated using the difference in RT/accuracy between high-probability and low-probability triplets. At the same time, visuomotor performance was operationalized by overall task RT/accuracy and its changes over time, indicated by faster/less accurate responses in later blocks, regardless of triplet probability. Initially, we assessed implicit probabilistic learning and its association with OC tendencies during the experiment’s Learning phase.

Trial-level RT (log-transformed) and accuracy (coded as 0 for incorrect and 1 for correct) served as the outcome variables. We utilized Linear Mixed Models (LMMs) for analyzing RT, and Generalized Linear Mixed Models (GLMMs) for analyzing accuracy. Both LMMs and GLMMs were fit with the *mixed* function from the *afex* package ((56). These models included fixed effects for Time (In Study 1: Factor: Block 1-5, Block 6-10, Block 11-15, Block 16-20, Block 21-25 and in Study 2 as Factor: Block 1-5, Block 6-10, Block 11-15), Triplet Type (Factor: High, Low), OC tendencies (OCI-R score), and their higher-order interactions. OCI-R score was a continuous variable, therefore, it was mean-centered. Please note that, although we used the OCI-R score as a continuous variable for all our analyses, for more illustrative data visualization, we depicted results at lower OC tendencies (Mean OCI-R score - SD) and higher OC tendencies (Mean OCI-R score + SD), based on standard deviations from the mean. The models also included participant-specific intercepts and correlated slopes for the within-subject variables of Time and Triplet Type, treated as random effects. In reporting the results, the main effects of Time is interpreted as improvement in visuomotor performance both on RT and accuracy, regardless of probabilistic patterns. Meanwhile, the main effect of Triplet Type is seen as an indicator of the amount of implicit probabilistic learning, and the interaction between Time and Triplet Type represents the temporal trajectory of probabilistic learning throughout the task. In the learning phase, the last level of the Time factor was always the reference: in Study 1, it was Block 21-25, whereas in Study 2, it was Block 11-15.

Hence, our models were structured according to the following general form in lmer syntax:

Reaction time: *RT ∼ Time * Triplet Type * OCI-R score + (Time + Triplet Type* | *Participant)*

Accuracy: *Accuracy ∼ Time * Triplet Type * OCI-R score + (Time + Triplet Type* | *Participant)*

##### Interference phase

To evaluate updating, we incorporated an interference phase within the task, as described in the “Alternating Serial Reaction Time task” section. In the Interference phase of Study 2, there was a change in the ASRT task structure. In Block 20-25, participants got the reversed version of the original sequence, which also led to some changes in the predictability of the triplets. In the Interference phase, we investigated two forms of learning: Old knowledge and New knowledge. Probabilistic learning is computed by comparing RT or accuracy between trials with changed occurrence probabilities (’H L’ or ‘L H’) and trials with unchanged occurrence probabilities (’L L’). The difference between ‘H L’ and ‘L L’ indicated the retention of previously learned knowledge (or Old knowledge), while the difference in RT or accuracy between ‘L H’ and ‘L L’ indicated the acquisition of New knowledge. This meant that participants were expected to show increasingly faster responses/higher accuracy in ‘L H’ and ‘H L’ trials compared to ‘L L’ trials in Block 20-25. ‘L L’ trials served as a baseline for these learning scores, controlling for general practice effects. Since ‘L L’ trials were consistently low-probability across phases, no significant increase in accuracy was expected due to probability-based learning.

Similarly to the analysis of the Learning phase, Trial-level RT (log-transformed) and accuracy (coded as 0 for incorrect and 1 for correct) served as the outcome variables for the updating analysis. We utilized LMMs for analyzing RT, and GLMMs for analyzing accuracy. Models included fixed effects for Time (Factor: Block 16-20, Block 21-25, Block 26-30), Triplet Type (Factor: For ‘Old knowledge’‘H L’, ‘L L’, For ‘New knowledge’‘L H’, ‘L L’,), OC tendencies (mean-centered OCI-R score), and their higher-order interactions. The models also included participant-specific intercepts and correlated slopes for the within-subject variables of Time and Triplet Type, treated as random effects. In reporting the results, the main effect of Time is interpreted as a change in visuomotor performance both on RT and accuracy, regardless of probabilistic patterns. While the main effect of Triplet Type is seen as an indicator of the amount of implicit probabilistic learning, the interaction between Time and Triplet Type represents the temporal trajectory of learning. In the interference phase, the last level of the Time factar was always the reference (Block 26-30). Please note that, for all our analyses, we used the OCI-R score as a continuous variable, but for more illustrative data visualization, we depicted results at lower OC tendencies (Mean OCI-R score - SD) and higher OC tendencies (Mean OCI-R score + SD), based on standard deviations from the mean.

For all analyses, we evaluated assumptions of linearity, homoscedasticity, and normality of residuals using scatterplots and QQ plots of residuals. The assumptions of linearity and homoscedasticity were satisfied for all models. However, some models exhibited non-normal residuals. Nonetheless, fixed effects estimates of LMMs and GLMMs are known to be highly robust against this particular assumption violation (57). Therefore, we considered our models to be adequate. Sample sizes, including the total number of data points and sampling units, random effects estimates, Nakagawa’s marginal and conditional R^2^ (58) and adjusted Intraclass Correlation Coefficients (ICCs) are consistently reported in the summary tables (see in Supplementary Materials S2-S9). Post-hoc contrasts were performed using the *emmeans* R package after regridding (59). For fixed effects inference, we utilized Type III tests (for RT models) and Likelihood Ratio Tests (for accuracy models) by comparing nested models with and without the effect of interest. Figures were generated with *ggplot2* (60). P values from all post-hoc tests were adjusted for multiple comparisons using the Šidák method (61). Alpha level .05 was applied to all analyses, and all p-values are two-tailed. All data and code required to reproduce the results of this study are available on the OSF repository (https://osf.io/fae2z/)

## Results

### How did obsessive-compulsive tendencies relate to probabilistic learning? - Study 1

The RT model indicated a statistically significant main effect of Time (F_(4,172.93)_ = 91.55, *p* < .001), showing that participants’ response times improved in subsequent blocks (Block 1-5: 375 ms, 95% CI = [369, 381]; Block 6-10: 371 ms, 95% CI = [366, 377]; Block 11-15: 363 ms, 95% CI = [358, 368]; Block 16-20: 355 ms, 95% CI = [350, 359]; Block 21-25: 349 ms, 95% CI = [344, 354]; (Block 1-5) - (Block 6-10) contrast *p* < .005, (Block 1-5) - (Block 11-15) contrast *p* < .001, (Block 1-5) - (Block 16-20) contrast *p* < .001, (Block 1-5) -(Block 21-25) contrast *p* < .001, (Block 6-10) - (Block 11-15) contrast *p* < .001, (Block 6-10) - (Block 16-20) contrast *p* < .001; (Block 6-10) - (Block 21-25) contrast *p* < .001, (Block 11-15) - (Block 16-20) contrast *p* < .001, (Block 11-15) - (Block 21-25) contrast *p* < .001; (Block 16-20) - (Block 21-25) contrast *p* < .001). There was a statistically significant main effect of Triplet Type (F_(1,159.69)_ = 306.70, *p* < .001), with faster RTs for high-probability triplets than for low-probability triplets, indicating implicit probabilistic learning (High: 359 ms, 95% CI = [354, 364]; Low: 366 ms, 95% CI = [362, 371]). A statistically significant Time × Triplet Type interaction was observed (F_(4, 243525.33)_ = 20.83, *p* < .001), which reflected an enhancement in implicit probabilistic learning as time progressed (Low – High contrast in Block 1-5: 3.59 ms, 95% CI = [2.13, 5.05]; in Block 6-10: 5.98 ms, 95% CI = [4.52, 7.43]; in Block 11-15: 8.16 ms, 95% CI = [6.74, 9.58]; in Block 16-20: 10.14 ms, 95% CI = [8.74, 11.53]; in Block 21-25: 9.95 ms, 95% CI = [8.86, 11.33]; (Block 1-5) - (Block 6-10) contrast *p* < .113, (Block 1-5) - (Block 11-15) contrast *p* < .001, (Block 1-5) - (Block 16-20) contrast *p* < .001, (Block 1-5) -(Block 21-25) contrast *p* < .001, (Block 6-10) - (Block 11-15) contrast *p* < .186, (Block 6-10) - (Block 16-20) contrast *p* < .001; (Block 6-10) - (Block 21-25) contrast *p* < .001, (Block 11-15) - (Block 16-20) contrast *p* < .227, (Block 11-15) - (Block 21-25) contrast *p* < .412; (Block 16-20) - (Block 21-25) contrast *p* > .999). The main effect of the OCI-R score was not significant (F_(1,161.99)_ = 0.95, *p* = .330), indicating that OC tendencies were not associated withe baseline reaction time. Changes in visuomotor performance were also intact along the OC- spectrum as presented by the absence of a statistically significant Time × OCI-R interaction (F_(4, 172.68)_ = 1.45, *p* = .220). Additionally, no significant interaction was found for either the Triplet Type × OCI-R score (F_(1, 159.45)_ = 0.11, *p* < .736) and Triplet Type × Time × OCI-R score (F_(4, 243622.68)_ = 1.66, *p* = .157), which suggests that OC tendencies did not influence the amount and trajectory of implicit probabilistic learning (Fig. 4A).The full list of fixed and random effect parameters can be found in Supplementary Table S2.

**Figure 4.**
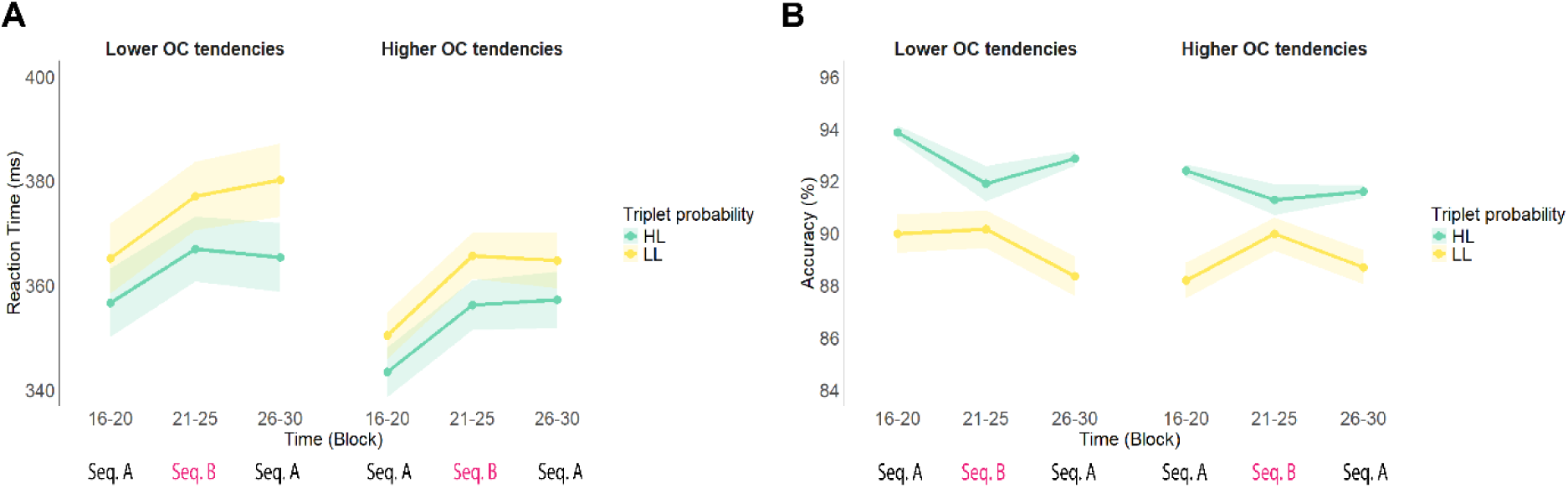
Participants’ performance during the Interference phase of Study 2 in terms of Old knowledge. **A. Reaction time**. The green line represents the responses to ‘H L’ probability triplets and the yellow line represents the responses to ‘L L’ probability triplets. The x-axis indicates Time (Block 16-20, Block 21-25, Block 26-30), the y-axis indicates mean reaction time (in ms). Triplets are now categorized based on the changes in their predictability between Sequence A and Sequence B. Participants showed a resistance of previously acquired implicit knowledge to interference as they retained the performance difference between ‘H L’ and ‘L L’ triplets even after exposure to Sequence B. **B. Accuracy**. The green line represents the responses given to the ‘H L’ probability and the yellow line represents the responses given to the ‘L L’ probability triplets. The x-axis indicates Times (4-6) y-axis indicates mean accuracy (in %). Triplets are now categorized based on the changes in their predictability between Sequence A and Sequence B. Participants showed a resistance of previously acquired implicit knowledge to interference as they retained the performance difference between ‘H L’ and ‘L L’ triplets even after exposure to Sequence B in terms of accuracy. Note that, OC tendencies was treated as a continuous variable in our analyses, but for more illustrative data visualization, we exhibited results at Lower OC tendencies (OCI-R:=13.93 −11.49, on the left) and Higher OC tendencies OCI-R:= 13.93 +11.49, on the right) based on standard deviations from the mean. In all subfigures, points indicate means and error bars indicate SEM.

**Figure 5.**
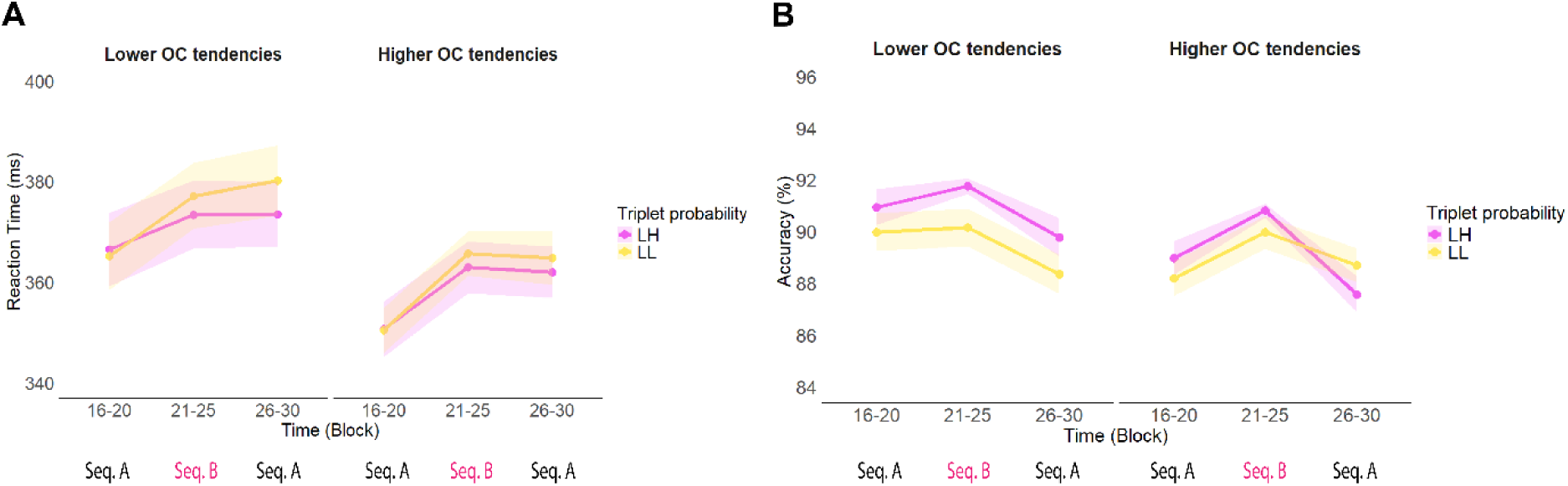
Participants’ performance during the Interference phase of Study 2 in terms of New knowledge. **A. Reaction time.** The pink line represents the responses given to the ‘L H’ probability triplets and the yellow line represents the responses given to the ‘L L’ probability triplets. The X-axis indicates Time (4-6) y-axis indicates mean reaction time (in ms). Triplets are now categorized based on the changes in their predictability between Sequence A and Sequence B. Participants showed a successful acquisition of novel implicit knowledge as they started to differentiate between ‘L H’ and ‘L L’ triplets in Sequence B, however, it was not modulated by the level of the OC tendencies. **B. Accuracy.** The pink line represents the responses given to the ‘ L H’ probability triplets and the yellow line represents the responses given to the ‘L L’ probability triplets. The X-axis indicates Time (4-6) y-axis indicates mean accuracy (in %). Participants demonstrated a successful acquisition of novel implicit knowledge as they started to differentiate between‘ L H’ and ‘L L’ triplets in Sequence B, however, it was not modulated by the severity of the OC tendencies. Note that, OC tendencies was treated as a continuous variable in our analyses, but for more illustrative data visualization, we exhibited results at Lower OC tendencies (OCI-R:=13.93 −11.49, on the left) and Higher OC tendencies (OCI-R:= 13.93 +11.49, on the right) based on standard deviations from the mean. In all subfigures, points indicate means and error bars indicate SEM.

With regards to accuracy, the model resulted in a statistically significant main effect of Time (χ^2^ (4) = 40.07, *p* < .001), indicating that participants became less accurate in subsequent blocks (Block 1-5: 92.1%, 95% CI = [91.4, 92.7]; Block 6-10: 91.2 %, 95% CI = [90.4, 91.8]; Block 11-15: 91.1 %, 95% CI = [90.5, 91.8]; Block 16-20: 90.6 %, 95% CI = [89.9, 91.2]; Block 20-25: 90.0 %, 95% CI = [89.2, 90.7]; (Block 1-5) - (Block 6-10) contrast *p* = .002, (Block 1-5) - (Block 11-15) contrast *p* = .009, (Block 1-5) - (Block 16-20) contrast *p* < .001, (Block 1-5) - (Block 21-25) contrast *p* < .001, (Block 6-10) - (Block 11-15) contrast *p* > .999, (Block 6-10) - (Block 16-20) contrast *p* = .296; (Block 6-10) - (Block 21-25) contrast *p* = .003, (Block 11-15) - (Block 16-20) contrast *p* = .177, (Block 11-15) - (Block 21-25) contrast *p* < .001; (Block 16-20) - (Block 21-25) contrast *p* = .136). There was also a statistically significant main effect of Triplet Type (χ^2^ (1) = 131.58, *p* < .001), with higher accuracy for high-probability triplets than for low-probability triplets, demonstrating implicit probabilistic learning (High: 92.3%, 95% CI = [91.7, 92.8]; Low: 89.6%, 95% CI = [88.9, 90.3]). A statistically significant Time × Triplet Type interaction showed an improvement in probabilistic learning as the task progressed (χ^2^ (4) = 21.89, *p* < .001) (High – Low contrast in Block 1-5: 1.50%, 95% CI = [0.93, 2.06]; in Block 6-10: 2.60%, 95% CI = [1.97, 3.23]; in Block 11-15 2.60%, 95% CI = [1.98, 3.22]; Block 16-20: 3.33%, 95% CI = [2.68, 3.98]; in Block 21-25: 3.56%, 95% CI = [2.88, 4.24]; (Block 1-5) - (Block 6-10) contrast *p* = .032, (Block 1-5) - (Block 11-15) contrast *p* = .032, (Block 1-5) - (Block 16-20) contrast *p* < .001, (Block 1-5) - (Block 21-25) contrast *p* < .001, (Block 6-10) - (Block 11-15) contrast *p* > .999, (Block 6-10) - (Block 16-20) contrast *p* = .493; (Block 6-10) - (Block 21-25) contrast *p* = .164, (Block 11-15) - (Block 16-20) contrast *p* = .503, (Block 11-15) - (Block 21-25) contrast *p* = .167; (Block 16-20) - (Block 21-25) contrast *p* > .999). The main effect of OCI-R score revealed that higher OC tendencies were associated with more accurate responses (χ^2^ (1) = 5.08, *p* = .024) Low OC tendencies: 90.3%, 95% CI = [89.4, 91.1]; High OC tendencies: 91.7%, 95% CI = [90.9, 92.4]. Similarly to the RT model, neither interaction involving OCI-R score was statistically significant, indicating that OC tendencies did not substantially influence probabilistic learning or its dynamics as assessed by accuracy (Triplet Type × OCI-R score: χ^2^ (1) = 0.27, *p* = .605; Time × Triplet Type × OCI-R score: χ^2^ (4) = 6.11, *p* = .191) as seen Fig. 4C. The full list of fixed and random effect parameters can be found in Supplementary Table S3.

To investigate the relationship between OC tendencies and probabilistic learning, we examined the correlation between the OCI-R total score and the difference learning score. The difference score was calculated as the difference in median RT and mean accuracy between high- and low-probability trials across the task’s temporal trajectory (Block 1–5, Block 6–10, Block 11–15, Block 16–20, Block 21–25), and then averaged across these five segments. Neither the correlation between probabilistic learning and OC tendencies in RT (*r*(162) = -.029, *p* = .709) nor in accuracy (*r*(162) = -.035; *p* = .653) was significant (Fig 4 C, D).

#### Summary of Study 1

In Study 1, we examined the relationship between the acquisition of probability-based structures and OC tendencies in the general population. We assessed implicit probabilistic learning with an unsupervised learning paradigm. Our results indicate that OC tendencies are not associated with individual differences in the acquisition of implicit probabilistic information, as presented by the lack of significant interactions involving OCI-R score during our analyses. Overall, this result implies that probabilistic learning is unaffected along the spectrum of subclinical OC tendencies. Nevertheless, the complexity of learning renders it even more interesting to examine the updating ability of learned probabilistic representations, as behaviour is strongly influenced not only by the information learned but also by the ability to manipulate it, especially in terms of OC behavioural representations. Thus, in order to gain a deeper understanding of maladaptive behavioural patterns in OCD, in Study 2, we were interested in both the resistance of acquired predictive representations and their updating ability to a new condition among OC tendencies.

#### Results of Study 2

To evaluate the stability of the previously acquired information, in the Interference phase of Study 2, the task structure was changed. In Block 21-25, unbeknownst to the participants, we introduced the reversed version of the original sequence. For the analysis of this phase, we categorized triplets based on their predictability in the two sequences: ‘H L’ triplets had high-probability in the original sequence, but low-probability in the new sequence, ‘L H’ triplets had low-probability in the original sequence, but high-probability in the new sequence, and ‘L L’ triplets were low-probability in both, serving as a baseline during our analysis. Previously established knowledge (Old knowledge) is represented by the performance difference between ‘L L’ and ‘H L’ triplets, while the acquisition of new probabilistic information (New knowledge) is measured by comparing performance between ‘L L’ and ‘L H’ triplets.

To explore how OC tendencies affected the resistance to previously acquired knowledge (Old knowledge) and the adjustment to novel information (New knowledge), we used trial-level (log-transformed) RT and accuracy during the Interference phase as outcome variables. The fixed effects in the models included Time factor (Block 15-20, Block 21-25, Block 25-30), Triplet Type (for Old knowledge: ‘H L’, ‘L L’, and for New knowledge: ‘L H’ and ‘L L’), OCI-R score (as a continuous variable), and their interactions. Note that the main effect of Triplet Type and its interactions indicate differences in learning, while main effects and interactions excluding this factor can be interpreted as differences in visuomotor performance. Furthermore, the OCI-R score was treated as a continuous variable in our analyses. However, for more illustrative data visualization, we presented results at Lower OC tendencies (M-SD) and Higher OC tendencies (M+SD) based on standard deviations from the mean.

### How did obsessive-compulsive tendencies modulate the resistance to previously acquired probabilistic information? - Old knowledge

In order to investigate the role of OC tendencies in updating predictive representations, we first also report performance in the Learning phase for both RT and accuracy. Then, we can properly interpret resistance to the acquired information. In terms of RT, the model yielded a statistically significant main effect of Time (F_(2,273.09)_ = 44.87, *p* < .001), showing that participants became faster in subsequent Blocks (Block 1-5: 384 ms, 95% CI = [380, 389]; Block 6-10: 384 ms, 95% CI = [380, 388]; Block 11-15: 376 ms, 95% CI = [372, 380]; (Block 1-5) - (Block 6-10) contrast *p* = .763, (Block 1-5) - (Block 11-15) contrast *p* < .001, (Block 6-10) - (Block 11-15) contrast *p* < .001). There was a statistically significant main effect of Triplet Type (F_(1,250.99)_ = 183.17, *p* < .001), with faster reaction times for high-probability triplets than for low-probability triplets, indicating implicit probabilistic learning (High: 379 ms, 95% CI = [375, 383]; Low: 384 ms, 95% CI = [380, 388]). A statistically significant Time × Triplet Type interaction was observed (F_(2, 227205.52)_ = 11.51, *p* < .001), which reflected a temporal enhancement in implicit probabilistic learning (Low – High contrast in Block 1-5: 3.97 ms, 95% CI = [2.76, 5.17]; in Block 6-10: 5.06 ms, 95% CI = [3.85, 6.27]; in Block 11-15: 7.51 ms, 95% CI = [6.33, 8.70]; (Block 1-5) -(Block 6-10) contrast *p* = .425, (Block 1-5) - (Block 11-15) contrast *p* < .001, (Block 6-10) - (Block 11-15) contrast *p* = .006). Neither the main effect nor any interactions involving OCI-R score were statistically significant, implying that OC tendencies did not influence implicit probabilistic learning. The full list of fixed and random effect parameters can be found in Supplementary Table S4.

In terms of Old knowledge, the RT model produced a statistically significant main effect of Time (F_(2,381.26)_ = 103.63, *p* < .001). Participants became generally slower after being exposed to the new sequence during Block 21-25 (Block 16-20: 350 ms, 95% CI = [346, 354]; Block 21-25: 361 ms, 95% CI = [358, 365]; Block 26-30: 363 ms, 95% CI = [359, 367]; (Block 16-20) - (Block 21-25) contrast *p* < .001, (Block 21-25) - (Block 26-30) contrast *p* < .001, (Block 21-25) - (Block 26-30) contrast *p* = .081). There was a statistically significant main effect of Triplet Type (F_(1, 270.98)_ = 249.67, *p* < .001), with faster reaction times to ‘H L’ triplets than to ‘L L’ triplets (‘H L’: 354 ms, 95% CI = [350, 358]; ‘L L’: 362 ms, 95% CI = [359, 366]). This suggests that, overall, participants maintained implicit knowledge of the old sequence despite being exposed to the new sequence. Additionally, the absence of a statistically significant Time × Triplet Type interaction (F_(2, 156913.57)_ = 1.86 *p* = .156) indicates that this resistance to interference was consistent across all Blocks. However, this resistance was not modulated by OC tendencies (Fig. 4A), based on the absence of statistically significant interactions involving OCI-R score: (Time x OCI-R score: F_(2, 281.95)_ = 2.57, *p* = .079; Triplet Type × OCI-R score: F_(1, 272.09)_ = 1.85, *p* = .175; Time × Triplet Type × OCI-R score: F_(2, 156937.86)_ = 1.42, *p* = .242), as well as main effect of OCI-R score F_(1, 255.25)_ = 0.81, *p* = .370). The full list of fixed and random effect parameters can be found in Supplementary Table S5.

For the accuracy indices, we first also report participants’s performance in the Learning phase. Similar to RT, the accuracy model resulted in a statistically significant main effect of Time (χ^2^ (2) = 56.61, *p* < .001), showing that participants became less accurate in later blocks (Block 1-5: 92.8%, 95% CI = [92.3, 93.3]; Block 6-10: 91.4 %, 95% CI = [90.9, 91.9]; Block 11-15: 91.1 %, 95% CI = [90.6, 91.6]; (Block 1-5) - (Block 6-10) contrast *p* < .001, (Block 1-5) - (Block 11-15) contrast *p* < .001, (Block 6-10) - (Block 11-15) contrast *p* = .263). There was a statistically significant main effect of Triplet Type (χ^2^ (1) = 161.25, *p* < .002), with higher accuracy for high-probability triplets than for low-probability triplets, evidencing implicit probabilistic learning (High: 92.8%, 95% CI = [92.4, 93.2]; Low: 90.7%, 95% CI = [90.2, 91.2]). Moreover, Time × Triplet Type showed a significant interaction, indicating improved probabilistic learning with time (χ^2^ (2) = 22.96, *p* < .001), (High – Low contrast in Block 1-5: 1.35%, 95% CI = [0.94, 1.76]; in Block 6-10: 2.12%, 95% CI = [1.67, 2.58]; in Block 11-15: 3.13%, 95% CI = [2.65, 3.60]; (Block 1-5) - (Block 6-10) contrast *p* = .246, (Block 1-5) - (Block 11-15) contrast *p* < .001, (Block 6-10) - (Block 11-15) contrast *p* = .004). The main effect of OCI-R score indicated less accurate responses with a higher level of OC tendencies (χ^2^ (1)= 6.43, *p* = .011; Low OC tendencies: 92.3%, 95% CI = [91.8, 92.9]; High OC tendencies: 91.2%, 95% CI = [90.6, 91.9]). However, in terms of visuomotor performance improvement, OC tendencies had no influence (Time × OCI-R score: χ^2^ (2) = 0.65*, p* = .724). There was no interaction between Triplet Type × OCI-R score (χ^2^(1) = 0.22, *p* = .641), showing that the prevalence of OC tendencies did not influence the amount of implicit probabilistic learning. Meanwhile, a statistically significant Time × Triplet Type × OCI-R score interaction revealed that participants with higher OC tendencies demonstrated faster extraction of probability-based structures (χ^2^ (2) = 7.07, *p* = .029) as assessed by accuracy: Lower OC tendencies: Block 1-5: 1.37%, 95% CI = [0.82, 1.92]; in Block 6-10: 1.69%, 95% CI = [1.08, 2.30]; in Block 11-15: 3.32%, 95% CI = [2.68, 3.96]; (Block 1-5) - (Block 6-10) contrast *p* = .805, (Block 1-5) - (Block 11-15) contrast *p* < .001, (Block 6-10) - (Block 11-15) contrast *p* < .001). Higher OC tendencies: Block 1-5: 1.32%, 95% CI = [0.73, 1.90]; in Block 6-10: 2.60 %, 95% CI = [1.95, 3.26]; in Block 11-15: 2.89 %, 95% CI = [2.21, 3.57]; Block 1-2 contrast *p* = .007, Block 1-3 contrast *p* < .001, Block 2-3 contrast *p* = .893). The full list of fixed and random effect parameters can be found in Supplementary Table S6.

The accuracy model regarding Old knowledge resulted in a statistically significant main effect of Time (χ^2^ (2) = 11.74, *p* = .003): participants became less accurate in the upcoming blocks (Block 16-20: 92.1%, 95% CI = [91.6, 92.5]; Block 21-25: 91.4 %, 95% CI = [90.9, 91.9]; Block 26-30: 91.3 %, 95% CI = [90.8, 91.8]; (Block 16-20) - (Block 21-25) contrast *p* = .021, (Block 16-30) - (Block 26-30) contrast *p* = .006, (Block 21-25) - (Block 26-30) contrast *p* = .986). There was a statistically significant main effect of Triplet Type (χ^2^ (1) = 108.03, *p* < .001), with higher accuracy for ‘H L’ trials than for ‘L L’ trials, indicating the resistance of the previously acquired information (‘H L’: 92.8%, 95% CI = [92.4, 93.2]; ‘L L’: 90.2%, 95% CI = [89.7, 90.7]). There was also a statistically significant Time × Triplet Type interaction (χ^2^ (2) = 21.68, *p* < .001), which stemmed from increasing probabilistic learning with the progression of the task (‘H L’ –’ L L’ contrast in Block 16-20: 3.23%, 95% CI = [2.60, 3.85]; in Block 21- 25: 1.39%, 95% CI = [0.64, 2.14]; in Block 26-30: 3.01%, 95% CI = [2.36, 3.65]; (Block 16-20) - (Block 21-25) contrast *p* < .001, (Block 16-20) - (Block 26-30) contrast *p* = .928, (Block 21-25) - (Block 26-30) contrast *p* = .001). OC tendencies had no influence on the accuracy of responses in the interference phase (OCI-R score main effect: χ^2^ (1) = 0.87, *p* = .350), nor on the improvement of visuomotor performance (Time × OCI-R score: χ^2^ (2) = 2.07, *p* = .355). The absence of Triplet Type × OCI-R score interaction indicates that OC tendencies did not modulate the resistance to previously acquired probabilistic information (Triplet Type × OCI-R score: χ^2^ (1) = 0.18, *p* = .673), and the progression of learning (Time x Triplet Type × OCI-R score: χ^2^ (2) = 1.40, *p* = .496) as assessed by accuracy (Fig. 4B).The full list of fixed and random effect parameters can be found in Supplementary Table S7.

### How did obsessive-compulsive tendencies relate to the ability to acquire novel probabilistic information? - New knowledge

The RT model effectively identified the impact of exposure to the new sequence, leading to a statistically significant main effect of Time (F_(2, 269.05)_ = 98.92, *p* < .001), with participants becoming overall slower after the introduction of the new sequence in Block 21-25 (Block 16- 20: 354 ms, 95% CI = [350, 358]; Block 21-25: 364 ms, 95% CI = [360, 368]; Block 26-30: 367 ms, 95% CI = [363, 370]; (Block 16-20) - (Block 21-25) contrast *p* < .001, (Block 16-20) - (Block 26-30) contrast *p* < .001, (Block 21-25) - (Block 26-30) contrast *p* = .004). There was a statistically significant main effect of Triplet Type (F_(1,274.79)_ = 12.26, p < .001), with faster reaction times for ‘L H’ triplets than for ‘L L’ triplets (‘L H’: 360 ms, 95% CI = [357, 364]; ‘L L’: 362 ms, 95% CI = [359, 366]), which highlights that participants gained implicit knowledge of the new sequence. A statistically significant Time × Triplet Type interaction indicates the extraction of the new probabilistic information (F_(2, 110362.72)_ = 3.69, *p* = .025), (Low – High contrast in Block 16-20: 0.40 ms, 95% CI = [-1.37, 2.18]; in Block 21-25: 3.32 ms, 95% CI = [1.83, 4.81]; in Block 26-30: 2.56 ms, 95% CI = [0.71, 4.42]; (Block 16-20) - (Block 21-25) contrast *p* = .018, (Block 16-20) - (Block 26-30) contrast *p* = .199, (Block 21-25)- (Block 26-30) contrast *p* = .864). No main effect of OCI-R score (F_(1, 255.28)_ = 0.78, *p* = .379) or interaction involving OCI-R score was statistically significant, suggesting that OC tendencies did not modulate the acquisition of new implicit knowledge or visuomotor performance to interference during the interference phase of the experiment, as assessed by RT (Time × OCI-R score: F_(2, 270.94)_ = 2.94, *p* = .055; Triplet Type × OCI-R score: F_(1, 277.63)_ = 1.81, *p* = .180; Time × Triplet Type × OCI-R score: F_(2, 110358.50)_ = 1.20, *p* = .300).The full list of fixed and random effect parameters can be found in Supplementary Table S8.

In terms of accuracy, participants became less accurate in subsequent blocks as indicated by a main effect of Time (χ2 (2) = 39.09, *p* < .001). Block 16-20: 90.6%, 95% CI = [90.0, 91.2]; Block 21-25: 91.4 %, 95% CI = [90.9, 91.8]; Block 26-30: 89.6 %, 95% CI = [89.0, 90.2]; (Block 16-20) - (Block 21-25) contrast *p* = .032, (Block 16-20) - (Block 26-30) contrast *p* = .006, (Block 21-25) - (Block 26-30) contrast *p* < .001). There was a statistically significant main effect of Triplet Type (χ^2^ (1) = 6.58, *p* = .009), with higher accuracy for ‘L H’ trials than for ‘L L’ trials, indicating the acquisition of the novel implicit knowledge (’L H’: 90.9%, 95% CI = [90.4, 91.3]; ‘L L’: 90.3%, 95% CI = [89.8, 90.8]). There was also a statistically significant Time × Triplet Type interaction (χ^2^ (2) = 8.13, *p* = .017), which resulted from a better acquisition of the novel information with Time (’H L’ – ‘L L’ contrast in Block 16-20: 0.52%, 95% CI = [-0.13, 2.33]; in Block 21-25: 1.19%, 95% CI = [0.59, 1.78]; in Block 26-30: 0.94 %, 95% CI = [-0.81, 0.78]; Block 16-20 - Block 21-25 contrast *p* = .379, Block 16-20 - Block 26-30 contrast *p* = .691, Block 21-25 - Block 26-30 contrast *p* = .037). Similarly to RT, OC tendencies did not influence mean accuracy in the Interference phase, as shown by a lack of significant main effect of OCI-R score (χ^2^(1) = 1.65, *p* = .199). Furthermore, it indicated no modulating effect on visuomotor performance (Time × OCI-R score interaction: χ^2^ (2) = 0.60, *p* = .742), and nor did it influenc the ability to obtain new probabilistic information (Triplet Type × OCI-R score: χ^2^(1) = 1.29, *p* = .256), or its temporal progression (Time × Triplet Type × OCI-R score: χ^2^ (2) = 1.02, *p* = .600). The full list of fixed and random effect parameters can be found in Supplementary Table S9.

#### Summary of Study 2

In Study 2, we took a step further by investigating how OC tendencies influence individual differences in the general population regarding the resistance to and updating of predictive representations when adapting to volatile conditions. As in Study 1, we utilized the same unsupervised learning paradigm, the ASRT task, to assess these. To investigate the role of OC tendencies in updating predictive representations, it was fundamental to measure the acquisition of probability-based structures in Study 2, too. Our results demonstrated strong performance in learning; however, in terms of accuracy, higher OC tendencies were associated with faster extraction of probability-based structures. This suggests that while the amount of learning may not vary, individual differences in the pattern of learning could emerge along the spectrum of OC tendencies. In terms of updating, in contrast to prior studies (13,38,39,62), participants in the present experiment were not notified about any structural changes in the task. Our results indicate that OC tendencies are not associated with individual differences in the resistance of predictive representations, as evidenced by the lack of significant interactions involving OCI-R scores in the analysis related to ‘Old knowledge’. Similarly, we found no individual differences related to OC tendencies in the updating of predictive representations, as demonstrated by the results for ‘New knowledge’. Overall, this suggests that the resistance and updating of predictive representations are not influenced by OC tendencies in the general population.

## Discussion

In this study, we aimed to investigate potential individual differences in probabilistic learning and in updating predictive representations related to OC tendencies within the general population. We implemented online experiments on two independent non-clinical samples (N_Study1_ = 164, N_Study2_ = 257) measuring OC tendencies as a continuous spectrum from the mildest prevalence to the highest. In Study 1, we focused on the implicit unsupervised acquisition of probabilistic contingencies, and our results indicated that individuals with OC tendencies show robust implicit probabilistic learning. We took a step further in Study 2 by examining the ability to update predictive representations among individuals with OC tendencies. Our findings revealed that even individuals with high OC tendencies within a non-clinical population demonstrated superb updating capabilities of predictive representations. The results will be discussed here from three key perspectives: (I) the presence or absence of supervision, (II) the complexity or predictability of the information obtained and (III) the level of awareness during the task.

Initially, theories posited that obsessions and compulsions continuously reinforce each other to obtain a rewarding sense of relief or avoid perceived consequences. This proposition was supported by the majority of studies demonstrating intact supervised learning in individuals with OCD (13,17,18,23,37,62 but for exception see: 24,25), showing that supervision makes learning efficient in OCD. Nevertheless, learning can occur also in the absence of supervision, and this form of learning has yielded mostly weaker results in OCD research to date (27–32, but for exceptions see 33–35). Consequently, our findings corroborate previous studies, indicating that non-clinical individuals with different levels of OC tendencies are capable of learning effectively in an unsupervised context, just like individuals with clinical OCD (33–35). This suggests that, while supervised learning appears to be effective in individuals with OCD, inconsistencies persist in the context of unsupervised learning. These differences may reflect factors beyond the mere presence or absence of supervision in the learning process. Therefore, future research is needed to resolve these contradictions and explore additional factors that could lead to a more comprehensive understanding of learning in both OCD and non-clinical OC populations.

One such factor that might influence the learning process than supervision is the predictability and complexity of the obtained information. Our results add to our understanding of the acquisition of more complex, probabilistic contingencies in relation to OC tendencies and clinical OCD, demonstrating robust performance akin to previous studies (17,18,23 but for exception see: 24,25,38). Performance is also inconsistent when it comes to deterministic contingencies: easier associative patterns evoke efficient learning (13,14) while fixed, repetitive first or second-order adjacent patterns (i.e., where one or two subsequently presented stimuli predict the upcoming one) evoke both reduced performance (27–31) and intact performance in clinical OCD (34,35,37) and also along the spectrum of non-clinical OC tendencies (36,38). A recent study highlighted potential differences in performance under probabilistic versus deterministic conditions (38). Building on this, we have demonstrated that individuals with non-clinical OC tendencies can efficiently acquire higher-order probabilistic contingencies, specifically second-order non-adjacent ones. However, these findings alone are insufficient for drawing definitive conclusions. Further research is needed to elucidate the underlying factors influencing preferences for probabilistic or deterministic contingencies in both OCD and non-clinical OC populations.

A third key factor influencing learning processes in OCD and non-clinical OC populations might be the explicit awareness during acquisition. In most studies, participants were explicitly informed that they were engaged in a learning process, and these studies mostly demonstrated intact performance (13,14,31,36,37) In cases where participants were unaware that they were learning, studies have shown mixed results: some reported reduced performance (24,25,27–32) while others observed intact performance (17,18,23,33–35). Many of these studies have been criticized, as participants often acquired the tasks rapidly and efficiently, facilitating a transition from implicit to explicit learning. In fact, in some cases involving individuals with OCD, even superior performance was observed (27,28). Moreover, a study comparing explicit and implicit learning conditions found that individuals with OCD performed better when they had explicit knowledge of the learning process (31). Our current findings, however, contradict this interpretation, as our participants were able to efficiently acquire information without any awareness of the learning process—similarly to a recent study where learning could also remain entirely implicit (33). It is important to note that while this study did not directly assess the impact of task explicitness, we employed second-order, non-adjacent dependencies within a noisy environment, which clearly contributed to keeping the task entirely implicit (47,54). Future research is therefore needed to pinpoint the effect of explicit instructions on probabilistic contingencies, expanding on those applied to deterministic ones.

Taken together with all the empirical evidence discussed, several factors appear to influence learning processes in individuals with OCD and OC tendencies. Drawing specific conclusions about the role of supervision in learning remains challenging, as other elements— such as the complexity and predictability of the information, along with whether it is acquired implicitly or explicitly—also play crucial roles. In sum, it seems that these three factors collectively impact the learning processes, and it remains difficult to separate them at this point, as they dominate differently depending on the various methods of investigation. Nevertheless, our current findings contribute to the existing body of knowledge by incorporating all three factors in the same experimental design, and demonstrating the robust acquisition of probabilistic contingencies in an implicit, unsupervised manner among OC tendencies. Conducting similar studies in clinical OCD cases is highly warranted to distinguish how deterministic and probabilistic contingencies are acquired in both supervised and unsupervised manners, while also controlling for explicit and implicit awareness.

Beyond the implicit extraction of probabilistic contingencies, our study advances our knowledge by focusing on the updating of predictive representations among OC tendencies. Our results do not align with prior theories since the incidental updating of predictive representations remained superb among OC tendencies in a non-clinical population. Prominent theories of OCD highlight both an excessive bias toward well-established knowledge (13,14,37) and also difficulties in relying on previous knowledge (38,39). The incinsisteny between our results and the previous theories may stem from the fact that they measured supervised learning processes, in which reward-associated behaviour persisted for participants even after it had been devalued or diminished following explicit prior warnings (13,14,37). In our study, we did not provide warnings, hence, participants were unaware they are in a learning situation, and that the stimulus sequence structure changed during the task (from sequence A to sequence B and back to sequence A). This allowed us to capture a much more implicit, warning-free updating mechanism. On the other hand, previous studies focused on clinical OCD populations, so a potential difference in sample characteristics could also explain why our study could not confirm the findings.

Interestingly, our results do not align with the theory suggesting that individuals with subclinical OC tendencies rely less on past experiences and exhibit a preference for recent outcomes (38,39) as they showed suberb updating capabilities regarding both past experiences (Old knowledge) and new one (New knowledge). The theory was based on more explicit processes and the task did not involve excessive training necessary for the development of stable predictive representations (38). It was also posited that individuals face greater difficulties under probabilistic contingencies, leading to increased uncertainty and exploratory behaviour compared to deterministic conditions (38). Similar to this, previous research has also demonstrated that in deterministic situations, explicit awareness can enhance performance in individuals with OCD (31). Interestingly, they experienced difficulties in the probabilistic condition, which contradicts our current findings. This discrepancy could be explained, for instance, by the extended exposure to repetitive probabilistic patterns in our study, which may have helped participants overcome uncertainty and enabled the efficient updating of predictive representations without requiring explicit awareness in individuals with OC tendencies. In summary, existing theories on updating mechanisms in OCD suggest both a bias toward relying on past information and a reduced ability to effectively utilize it. Our approach offers a new perspective, and complements these two, by showing that individuals with subclinical OC tendencies can adapt to new situations without prior warning, while also retaining and incorporating new information. Future research is needed to further explore updating processes in OCD and OC tendencies.

This study has outlined the learning and updating mechanisms in individuals with OC tendencies while acknowledging that the maladaptive behavioural manifestations of OCD and OC tendencies can be shaped by the complex interplay of neurocognitive processes, specifically, between goal-directed and habitual (automatic) processes (13,14,19,37,63–65). Goal-directed systems encompass cognitive control functions that operate in a structured manner in order to support adaptive behaviour, regulate the achievement of desired goals, and the avoidance of undesired consequences (66,67). Although this theory is primarily based on a supervised, associative learning paradigm (13,14,37), it remains essential for interpreting our findings, as probabilistic learning also captures habitual processes, directly addressing a core aspect of habit formation (47). Hence, our results align with these findings, showing that probabilistic (or habitual) learning remained robust along the spectrum of subclinical OC tendencies in an unsupervised manner. At the same time, a theory based on probabilistic learning also emphasizes a dynamic interplay, highlighting the competitive relationship between habitual processes and the prefrontal cortex and its inherent goal-directed control processes (45,47,68–70). Even though this study did not focus directly on the interplay between neurocognitive processes, our findings on robust probabilistic learning lay the groundwork for bridging these two previously distinct research lines. To achieve this, future research is highly warranted to unravel how the interplay or imbalance between probabilistic learning and goal-directed control processes contributes to behavioural manifestations in both clinical OCD populations and non-clinical individuals with OC tendencies.

To the best of our knowledge, this is the first study to empirically test the unsupervised acquisition of probabilistic second-order non-adjacent dependencies and the updating of predictive representations among non-clinical individuals with OC tendencies in the general population. It went beyond the traditional dichotomous comparison between clinical and neurotypical groups by recruiting a large sample from the general population and accounting for individual differences in probabilistic learning processes. The major issue with dichotomous group comparisons is that OCD is a heterogeneous condition (71–75), with varying cognitive profiles also related to statistical learning (28,34). In contrast to our approach, such comparisons often limit the ability to uncover individual differences within small OCD samples. Examining a broad spectrum of individuals from the general population with OC tendencies enables a more nuanced analysis. However, this also complicates the interpretation of our results, as it remains uncertain whether there are qualitative differences between clinical and non-clinical populations. Following the example of Hoven et al. (76)’s metacognition study, researchers should next compare probabilitistic learning in individuals diagnosed with OCD and undiagnosed individuals who exhibit similar levels of OC tendencies. In the long term, this approach could help determine whether the relationship between clinical and non-clinical populations is linear or if there are substantial differences between the extremes of the spectrum and individuals with moderate symptom tendencies.

In summary, our study revealed robust probabilistic learning and superb updating capabilities of predictive representations associated with individual differences in OC tendencies in two large general population samples. Our results may provide a new direction for existing theories on implicit information processing in OCD and also along the spectrum of OC tendencies. Further multiprocess studies are necessary to understand the interaction of neurocognitive processes underlying the condition. This not only could contribute to a better understanding of OCD but could also enable the development of individualized prevention and therapeutic approaches in the future. Probabilistic learning might assist in forming and maintaining new skills and habits in psychotherapy. If this form of learning remains intact in OCD, we can build upon it to develop new procedures.One potential approach to alleviating the rigid, deterministic patterns and compulsive behaviors characteristic of OCD is the intentional incorporation of noise or probabilistic variability into processes. Such interventions aim to help individuals break free from compulsive behaviors by exposing them to uncertainty and enabling them to learn to accept it without resorting to ritualistic actions. Meanwhile, the results of this study showed that probabilistic learning, or in other words, skill or habit learning, remains superb in individuals with subclinical OC tendencies in the general population. Therefore, those who repeatedly check whether they have locked the front door or doubt that their wallet and phone are in their bag or pockets are still capable of extracting probability-based information from environmental cues and of properly predicting future events.

## Acknowledgements

This work was supported by the Chaire de Professeur Junior Program by INSERM and French National Grant Agency (ANR-22-CPJ1-0042-01) (to D.N.); the National Brain Research Program project NAP2022-I-2/2022 (to D.N.); Hungarian National Research, Development and Innovation Office Grant OTKA PD 146424 (to K.F.). We thank Gábor Csifcsák for his valuable feedback, which greatly contributed to the improvement of the work.

## Data availability statement

The data and analysis codes are available at the OSF repository (https://osf.io/fae2z/).

## Declaration of generative AI and AI-assisted technologies in the writing process

During the preparation of this work the author(s) used ChatGPT 3.5 to improve the writing style of our proposal. The author(s) reject using AI for scientific purposes to gain unethical benefits. However, the author(s) believe that it helps the equality of native and non-native English speakers to have the same opportunities. Thus, the author(s) do recommend using generative AI for lingual purposes (only). After using this tool/service, the author(s) reviewed and edited the content as needed and take(s) full responsibility for the content of the publication.

## Declaration of Competing Interest

The author(s) declare no competing interest.

## Notes

https://osf.io/fae2z/

### Competing Interest Statement

The authors have declared no competing interest.

