## Supplementary_materials for "Individual differences in probabilistic learning and updating predictive representations in individuals with obsessive-compulsive tendencies"

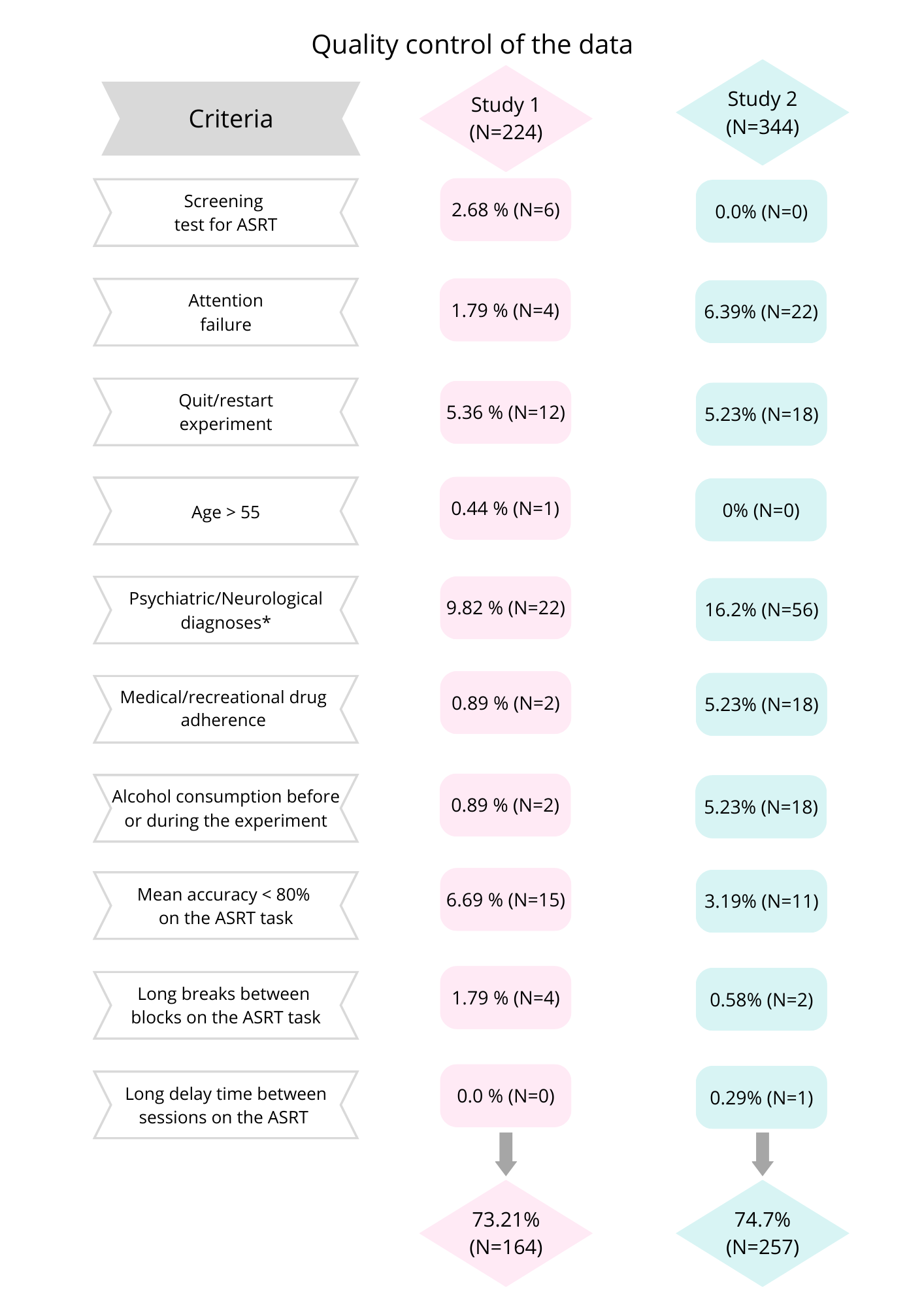

**Supplementary Figure S1.** Flow chart of quality control of the data. Since some participants met multiple exclusion criteria, a total of 60 participants were excluded from Study 1, resulting in a final sample comprising 73.21% of the initial participants. In Study 2, 87 participants were excluded, leaving 74.7% of the participants in the final sample.

*Based on self-reported neurological impairment, Epilepsy or head injury, or reported diagnosis of Autism Spectrum Disorder, Attention Deficit Hyperactivity Disorder, Obsessive-Compulsive Disorder, Schizophrenia or Psychotic Disorder, any type of Depressive Disorder, Anxiety Disorder, and/or Eating Disorder, as well as any Personality Disorder, and Post-Traumatic Stress Disorder were excluded from the analysis.

**Supplementary Table S2**. Results of the linear mixed model on log-transformed RT regarding probabilistic learning in Study 1.

|  | **Prediction of RT** | | | | | |
| --- | --- | --- | --- | --- | --- | --- |
| *Terms* | *b* | *SE* | *95% CI* | *t* | *df* | *p* |
| (Intercept) | 362.403 | 2.483 | 357.534 – 367.339 | 860.194 | 161.990 | **<0.001** |
| Block 1-5 [1] | 1.035 | 0.002 | 1.030 – 1.040 | 14.301 | 168.514 | **<0.001** |
| Block 6-10 [2] | 1.024 | 0.002 | 1.020 – 1.028 | 11.527 | 169.789 | **<0.001** |
| Block 11-15 [3] | 1.001 | 0.002 | 0.998 – 1.005 | 0.779 | 174.609 | 0.437 |
| Block 16-20 [4] | 0.979 | 0.002 | 0.975 – 0.982 | -11.537 | 174.930 | **<0.001** |
| Triplet Type [High] | 0.990 | 0.001 | 0.988 – 0.991 | -17.513 | 159.694 | **<0.001** |
| OCI-R score | 0.999 | 0.001 | 0.998 – 1.001 | -0.976 | 161.987 | 0.330 |
| Block 1-5 [1] x— Triplet Type [High] | 1.006 | 0.001 | 1.004 – 1.007 | 7.169 | 243491.238 | **<0.001** |
| Block 6-10 [2] x— Triplet Type [High] | 1.002 | 0.001 | 1.001 – 1.004 | 3.061 | 243497.789 | **0.002** |
| Block 11-15 [3] x— Triplet Type [High] | 0.999 | 0.001 | 0.998 – 1.001 | -0.886 | 243491.490 | 0.376 |
| Block 16-20 [4] x— Triplet Type [High] | 0.996 | 0.001 | 0.995 – 0.998 | -4.652 | 243500.591 | **<0.001** |
| Block 1-5 [1] x— OCI-R score | 1.000 | 0.000 | 1.000 – 1.000 | 0.140 | 168.494 | 0.889 |
| Block 6-10 [2] x— OCI-R score | 1.000 | 0.000 | 0.999 – 1.000 | -1.142 | 169.508 | 0.255 |
| Block 11-15 [3] x— OCI-R score | 1.000 | 0.000 | 1.000 – 1.000 | -1.070 | 174.532 | 0.286 |
| Block 16-20 [4] x— OCI-R score | 1.000 | 0.000 | 1.000 – 1.001 | 2.053 | 174.787 | **0.042** |
| Triplet Type [High] x OCI-R score | 1.000 | 0.000 | 1.000 – 1.000 | -0.338 | 159.451 | 0.736 |
| (Block 1-5 [1] x— Triplet Type [High] x — OCI-R score | 1.000 | 0.000 | 1.000 – 1.000 | -1.822 | 243478.403 | 0.068 |
| (Block 6-10 [2] x— Triplet Type [High] x — OCI-R score | 1.000 | 0.000 | 1.000 – 1.000 | 2.010 | 243500.616 | **0.044** |
| (Block 11-15 [3] x— Triplet Type [High] x — OCI-R score | 1.000 | 0.000 | 1.000 – 1.000 | 0.648 | 243484.789 | 0.517 |
| (Block 16-20 [4] x— Triplet Type [High] x — OCI-R score | 1.000 | 0.000 | 1.000 – 1.000 | -0.679 | 243494.478 | 0.497 |
| **Random Effects** | | | | | | |
| σ^2^ | 0.032 | | | | | |
| τ_00_ _Participant_ | 0.008 | | | | | |
| τ_11_ _Participant.Block 1-5_ | 0.001 | | | | | |
| τ_11_ _Participant.Block 6-10_ | 0.001 | | | | | |
| τ_11_ _Participant.Block 11-15_ | 0.000 | | | | | |
| τ_11_ _Participant.Block 15-20_ | 0.000 | | | | | |
| τ_11_ _Participant. Triplet Type [High]_ | 0.000 | | | | | |
| ρ_01_ | 0.209 | | | | | |
|  | 0.203 | | | | | |
|  | -0.133 | | | | | |
|  | -0.150 | | | | | |
|  | 0.393 | | | | | |
| ICC | 0.210 | | | | | |
| N _Participant_ | 164 | | | | | |
| Observations | 244258 | | | | | |
| Marginal R^2^ / Conditional R^2^ | 0.023 / 0.228 | | | | | |

**Note.** The marginal R-squared considers only the variance of the fixed effects, while the conditional R-squared takes both the fixed and random effects into account. Degrees of freedom are based on Satterthwaite’s approximation. Statistically significant terms are highlighted in bold. Terms in brackets indicate the level of factor that is contrasted against the reference level.

**Supplementary Table S3.** Results of the binomial generalized mixed model on accuracy regarding probabilistic learning in Study 1.

|  | **Prediction of Accuracy** | | | | | |
| --- | --- | --- | --- | --- | --- | --- |
| *Terms* | *Log-Odds* | *SE* | *95% CI* | *z* | *df* | *p* |
| (Intercept) | 2.314 | 0.036 | 2.243 – 2.385 | 63.864 | Inf | **<0.001** |
| Block 1-5 [1] | 0.137 | 0.025 | 0.089 – 0.186 | 5.544 | Inf | **<0.001** |
| Block 6-10 [2] | 0.018 | 0.021 | -0.024 – 0.060 | 0.843 | Inf | 0.399 |
| Block 11-15 [3] | 0.017 | 0.020 | -0.022 – 0.056 | 0.855 | Inf | 0.393 |
| Block 16-20 [4] | -0.052 | 0.019 | -0.090 – -0.014 | -2.687 | Inf | **0.007** |
| Triplet Type [High] | 0.163 | 0.011 | 0.141 – 0.185 | 14.418 | Inf | **<0.001** |
| OCI-R score | 0.007 | 0.003 | 0.001 – 0.014 | 2.272 | Inf | **0.023** |
| Block 1-5 [1] x— Triplet Type [High] | -0.061 | 0.015 | -0.090 – -0.032 | -4.106 | Inf | **<0.001** |
| Block 6-10 [2] x— Triplet Type [High] | -0.002 | 0.014 | -0.030 – 0.026 | -0.167 | Inf | 0.867 |
| Block 11-15 [3] x— Triplet Type [High] | -0.002 | 0.014 | -0.030 – 0.026 | -0.150 | Inf | 0.881 |
| Block 16-20 [4] x— Triplet Type [High] | 0.031 | 0.014 | 0.004 – 0.059 | 2.230 | Inf | **0.026** |
| Block 1-5 [1] x— OCI-R score | -0.003 | 0.002 | -0.007 – 0.002 | -1.170 | Inf | 0.242 |
| Block 6-10 [2] x— OCI-R score | 0.000 | 0.002 | -0.003 – 0.004 | 0.078 | Inf | 0.938 |
| Block 11-15 [3] x— OCI-R score | 0.000 | 0.002 | -0.003 – 0.004 | 0.165 | Inf | 0.869 |
| Block 16-20 [4] x— OCI-R score | 0.001 | 0.002 | -0.002 – 0.005 | 0.738 | Inf | 0.461 |
| Triplet Type [High] x OCI-R score | 0.001 | 0.001 | -0.001 – 0.002 | 0.518 | Inf | 0.605 |
| (Block 1-5 [1] x— Triplet Type [High] x — OCI-R score | 0.001 | 0.001 | -0.001 – 0.004 | 1.102 | Inf | 0.271 |
| (Block 6-10 [2] x— Triplet Type [High] x — OCI-R score | -0.002 | 0.001 | -0.004 – 0.001 | -1.356 | Inf | 0.175 |
| (Block 11-15 [3] x— Triplet Type [High] x — OCI-R score | -0.001 | 0.001 | -0.003 – 0.002 | -0.638 | Inf | 0.524 |
| (Block 16 -20 [4] x— Triplet Type [High] x — OCI-R score | -0.001 | 0.001 | -0.004 – 0.001 | -0.928 | Inf | 0.353 |
| **Random Effects** | | | | | | |
| σ^2^ | 3.290 | | | | | |
| τ_00_ _Participant_ | 0.208 | | | | | |
| τ_11_ _Participant.Block 1-5_ | 0.059 | | | | | |
| τ_11_ _Participant.Block 6-10_ | 0.036 | | | | | |
| τ_11_ _Participant.Block 11-15_ | 0.027 | | | | | |
| τ_11_ _Participant.Block 15-20_ | 0.025 | | | | | |
| τ_11_ _Participant. Triplet Type [High]_ | 0.011 | | | | | |
| ρ_01_ | 0.116 | | | | | |
|  | 0.185 | | | | | |
|  | -0.079 | | | | | |
|  | -0.255 | | | | | |
|  | 0.034 | | | | | |
| ICC | 0.073 | | | | | |
| N _Participant_ | 164 | | | | | |
| Observations | 269363 | | | | | |
| Marginal R^2^ / Conditional R^2^ | 0.010 / 0.082 | | | | | |

**Note.** The table shows regression coefficients of fixed effects and summary information about the random effects. Coefficients are log odds, thus positive values indicate that the given independent variable is associated with an increased likelihood of correct responses, and negative values are the opposite. P values for coefficients are from z tests, making the degrees of freedom infinite. The marginal R-squared considers only the variance of the fixed effects, while the conditional R-squared takes both the fixed and random effects into account. Statistically significant terms are highlighted in bold. Terms in brackets indicate the level of factor that is contrasted against the reference level.

**Supplementary Table S4.** Results of the linear mixed model on log-transformed RT regarding probabilistic learning in Study 2.

|  | **Prediction of RT** | | | | | |
| --- | --- | --- | --- | --- | --- | --- |
| *Terms* | *b* | *SE* | *95% CI* | *t* | *df* | *p* |
| (Intercept) | 381.301 | 2.026 | 377.332 – 385.313 | 1118.456 | 254.905 | **<0.001** |
| Block 1-5 [1] | 1.008 | 0.002 | 1.005 – 1.011 | 5.199 | 268.151 | **<0.001** |
| Block 6-10 [2] | 1.006 | 0.001 | 1.004 – 1.008 | 5.502 | 279.135 | **<0.001** |
| Triplet Type [High] | 0.993 | 0.001 | 0.992 – 0.994 | -13.534 | 250.990 | **<0.001** |
| OCI-R score | 1.000 | 0.000 | 0.999 – 1.000 | -0.882 | 254.944 | 0.379 |
| Block 1-5 [1] x — Triplet Type [High] | 1.002 | 0.001 | 1.001 – 1.003 | 3.508 | 227191.022 | **<0.001** |
| Block 6-10 [1] x — Triplet Type [High] | 1.001 | 0.001 | 0.999 – 1.002 | 1.090 | 227215.145 | 0.276 |
| Block 1-5 [1] x— OCI-R score | 1.000 | 0.000 | 1.000 – 1.000 | -1.026 | 268.791 | 0.306 |
| Block 6-10 [2] x— OCI-R score | 1.000 | 0.000 | 1.000 – 1.000 | 0.764 | 280.900 | 0.446 |
| Triplet Type [High] x— OCI-R score | 1.000 | 0.000 | 1.000 – 1.000 | 0.453 | 253.339 | 0.651 |
| (Block 1-5 [1] x— Triplet Type [High]) x— OCI-R | 1.000 | 0.000 | 1.000 – 1.000 | 1.310 | 227196.061 | 0.190 |
| (Block 6-10 [2] x— Triplet Type [High]) x— OCI-R | 1.000 | 0.000 | 1.000 – 1.000 | -0.813 | 227176.306 | 0.416 |
| **Random Effects** | | | | | | |
| σ^2^ | 0.033 | | | | | |
| τ_00_ _Participant_ | 0.007 | | | | | |
| τ_11_ _Participant. Block 1-5_ | 0.001 | | | | | |
| τ_11_ _Participant. Block 6-10_ | 0.000 | | | | | |
| τ_11_ _Participant.Triplet_Type[High]_ | 0.000 | | | | | |
| ρ_01_ | 0.144 | | | | | |
|  | 0.001 | | | | | |
|  | 0.110 | | | | | |
| ICC | 0.191 | | | | | |
| N _Participant_ | 257 | | | | | |
| Observations | 227889 | | | | | |
| Marginal R^2^ / Conditional R^2^ | 0.005 / 0.195 | | | | | |

**Note.** The marginal R-squared considers only the variance of the fixed effects, while the conditional R-squared takes both the fixed and random effects into account. Degrees of freedom are based on Satterthwaite’s approximation. Statistically significant terms are highlighted in bold. Terms in brackets indicate the level of factor that is contrasted against the reference level.

**Supplementary Table S5**. Results of the linear mixed model on log-transformed RT regarding ’Old knowledge’ in Study 2.

|  | **Prediction of RT** | | | | | |
| --- | --- | --- | --- | --- | --- | --- |
| *Terms* | *b* | *SE* | *95% CI* | *t* | *df* | *p* |
| (Intercept) | 358.244 | 1.887 | 354.547 – 361.979 | 1116.469 | 255.225 | **<0.001** |
| Block 16-20 [1] | 0.978 | 0.002 | 0.975 – 0.981 | -14.373 | 298.397 | **<0.001** |
| Block 21-25 [2] | 1.009 | 0.001 | 1.006 – 1.011 | 6.500 | 259.212 | **<0.001** |
| Triplet Type [H L] | 0.989 | 0.001 | 0.987 – 0.990 | -15.801 | 270.983 | **<0.001** |
| OCI-R score | 1.000 | 0.000 | 0.999 – 1.000 | -0.898 | 255.254 | 0.370 |
| Block 16-20 [1] x— Triplet Type [H L] | 1.001 | 0.001 | 1.000 – 1.003 | 1.570 | 156792.988 | 0.116 |
| Block 21-25 [2] x— Triplet Type [H L] | 1.000 | 0.001 | 0.998 – 1.002 | 0.018 | 156791.687 | 0.986 |
| Block 16-20 [1] x— OCI-R score | 1.000 | 0.000 | 0.999 – 1.000 | -1.793 | 299.581 | 0.074 |
| Block 21-25 [2] x— OCI-R score | 1.000 | 0.000 | 1.000 – 1.000 | 1.987 | 260.105 | **0.048** |
| Triplet Type [H L] x— OCI-R score | 1.000 | 0.000 | 1.000 – 1.000 | 1.359 | 272.087 | 0.175 |
| (Block 16-20 [1] x— Triplet Type [H L] x— OCI-R score | 1.000 | 0.000 | 1.000 – 1.000 | -0.534 | 156914.412 | 0.593 |
| (Block 21-25 [2] x— Triplet Type [H L] x— OCI-R score | 1.000 | 0.000 | 1.000 – 1.000 | -1.000 | 156723.370 | 0.317 |
| **Random Effects** | | | | | | |
| σ^2^ | 0.030 | | | | | |
| τ_00_ _Participant_ | 0.007 | | | | | |
| τ_11_ _Participant Block 16-20 [1]_ | 0.000 | | | | | |
| τ_11_ _Participant Block 21-25 [2]_ | 0.000 | | | | | |
| τ_11_ _Participant. Triplet Type [H L]_ | 0.000 | | | | | |
| ρ_01_ | 0.107 | | | | | |
|  | -0.046 | | | | | |
|  | 0.162 | | | | | |
| ICC | 0.205 | | | | | |
| N _Participant_ | 257 | | | | | |
| Observations | 157503 | | | | | |
| Marginal R^2^ / Conditional R^2^ | 0.011 / 0.214 | | | | | |

**Note.** The marginal R-squared considers only the variance of the fixed effects, while the conditional R-squared takes both the fixed and random effects into account. Degrees of freedom are based on Satterthwaite’s approximation. Statistically significant terms are highlighted in bold. Terms in brackets indicate the level of factor that is contrasted against the reference level.

**Supplementary Table S6.** Results of the binomial generalized mixed model on accuracy regarding probabilistic learning in Study 2.

|  | **Prediction of Accuracy** | | | | | |
| --- | --- | --- | --- | --- | --- | --- |
| *Terms* | *Log-Odds* | *SE* | *95% CI* | *z* | *df* | *p* |
| (Intercept) | 2.417 | 0.029 | 2.361 – 2.474 | 83.993 | Inf | **<0.001** |
| Block 1-5 [1] | 0.145 | 0.019 | 0.108 – 0.182 | 7.652 | Inf | **<0.001** |
| Block 6-10 [2] | -0.052 | 0.014 | -0.080 – -0.024 | -3.668 | Inf | **<0.001** |
| Triplet Type [High] | 0.143 | 0.010 | 0.124 – 0.161 | 14.997 | Inf | **<0.001** |
| OCI-R score | -0.006 | 0.002 | -0.011 – -0.001 | -2.555 | Inf | **0.011** |
| Block 1-5 [1] x — Triplet Type [High] | -0.042 | 0.011 | -0.064 – -0.019 | -3.672 | Inf | **<0.001** |
| Block 6-10 [1] x — Triplet Type [High] | -0.008 | 0.011 | -0.029 – 0.014 | -0.703 | Inf | 0.482 |
| Block 1-5 [1] x— OCI-R score | 0.000 | 0.002 | -0.003 – 0.003 | 0.024 | Inf | 0.981 |
| Block 6-10 [2] x— OCI-R score | 0.001 | 0.001 | -0.001 – 0.003 | 0.683 | Inf | 0.495 |
| Triplet Type [High] x— OCI-R score | -0.000 | 0.001 | -0.002 – 0.001 | -0.467 | Inf | 0.640 |
| (Block 1-5 [1] x— Triplet Type [High]) x— OCI-R | -0.000 | 0.001 | -0.002 – 0.001 | -0.371 | Inf | 0.710 |
| (Block 6-10 [2] x— Triplet Type [High]) x— OCI-R | 0.002 | 0.001 | 0.000 – 0.004 | 2.445 | Inf | **0.014** |
| **Random Effects** | | | | | | |
| σ^2^ | 3.290 | | | | | |
| τ_00_ _Participant_ | 0.194 | | | | | |
| τ_11_ _Participant. Block 1-5_ | 0.053 | | | | | |
| τ_11_ _Participant. Block 6-10_ | 0.017 | | | | | |
| τ_11_ _Participant.Triplet_Type[High]_ | 0.006 | | | | | |
| ρ_01_ | 0.351 | | | | | |
|  | -0.161 | | | | | |
|  | 0.448 | | | | | |
| ICC | 0.070 | | | | | |
| N _Participant_ | 257 | | | | | |
| Observations | 249134 | | | | | |
| Marginal R^2^ / Conditional R^2^ | 0.009 / 0.079 | | | | | |

**Note.** The table shows regression coefficients of fixed effects and summary information about the random effects. Coefficients are log odds, thus positive values indicate that the given independent variable is associated with an increased likelihood of correct responses, and negative values are the opposite. P values for coefficients are from z tests, making the degrees of freedom infinite. The marginal R-squared considers only the variance of the fixed effects, while the conditional R-squared takes both the fixed and random effects into account. Statistically significant terms are highlighted in bold. Terms in brackets indicate the level of factor that is contrasted against the reference level.

**Supplementary Table S7.** Results of the binomial mixed model on accuracy regarding ’Old knowledge’ in Study 2.

|  | **Prediction of Accuracy** | | | | | |
| --- | --- | --- | --- | --- | --- | --- |
| *Terms* | *Log-Odds* | *SE* | *95% CI* | *z* | *df* | *p* |
| (Intercept) | 2.389 | 0.026 | 2.339 – 2.440 | 93.505 | Inf | **<0.001** |
| Block 16-20 [1] | 0.062 | 0.018 | 0.026 – 0.098 | 3.414 | Inf | **0.001** |
| Block 21-25 [2] | -0.026 | 0.019 | -0.064 – 0.012 | -1.349 | Inf | 0.177 |
| Triplet Type [H L] | 0.166 | 0.014 | 0.138 – 0.193 | 11.774 | Inf | **<0.001** |
| OCI-R score | -0.002 | 0.002 | -0.006 – 0.002 | -0.938 | Inf | 0.348 |
| Block 16-20 [1] x— Triplet Type [H L] | 0.054 | 0.015 | 0.026 – 0.083 | 3.687 | Inf | **<0.001** |
| Block 21-25 [2] x— Triplet Type [H L] | -0.078 | 0.017 | -0.112 – -0.044 | -4.538 | Inf | **<0.001** |
| Block 16-20 [1] x— OCI-R score | -0.002 | 0.002 | -0.005 – 0.001 | -1.384 | Inf | 0.166 |
| Block 21-25 [2] x— OCI-R score | 0.002 | 0.002 | -0.001 – 0.005 | 1.036 | Inf | 0.300 |
| Triplet Type [H L] x— OCI-R score | -0.000 | 0.001 | -0.003 – 0.002 | -0.423 | Inf | 0.672 |
| (Block 16-20 [1] x— Triplet Type [H L] x—OCI-R score | 0.001 | 0.001 | -0.002 – 0.003 | 0.481 | Inf | 0.630 |
| (Block 21-25 [2] x— Triplet Type [H L] x—OCI-R score | 0.001 | 0.001 | -0.002 – 0.004 | 0.589 | Inf | 0.556 |
| **Random Effects** | | | | | | |
| σ^2^ | 3.290 | | | | | |
| τ_00_ _Participant_ | 0.133 | | | | | |
| τ_11_ _Participant Block 16-20 [1]_ | 0.024 | | | | | |
| τ_11_ _Participant Block 21-25 [2]_ | 0.012 | | | | | |
| τ_11_ _Participant. Triplet Type [H L]_ | 0.017 | | | | | |
| ρ_01_ | 0.159 | | | | | |
|  | -0.101 | | | | | |
|  | 0.108 | | | | | |
| ICC | 0.052 | | | | | |
| N _Participant.Public.ID_ | 257 | | | | | |
| Observations | 171403 | | | | | |
| Marginal R^2^ / Conditional R^2^ | 0.008 / 0.060 | | | | | |

**Note.** The table shows regression coefficients of fixed effects and summary information about the random effects. Coefficients are log odds, thus positive values indicate that the given independent variable is associated with an increased likelihood of correct responses, and negative values are the opposite. P values for coefficients are from z tests, making the degrees of freedom infinite. The marginal R-squared considers only the variance of the fixed effects, while the conditional R-squared takes both the fixed and random effects into account. Statistically significant terms are highlighted in bold. Terms in brackets indicate the level of factor that is contrasted against the reference level.

**Supplementary Table S8.** Results of the linear mixed model on log-transformed RT regarding ’New knowledge’ in Study 2.

|  | **Prediction of RT** | | | | | |
| --- | --- | --- | --- | --- | --- | --- |
| *Terms* | *b* | *SE* | *95% CI* | *t* | *df* | *p* |
| (Intercept) | 361.404 | 1.893 | 357.695 – 365.152 | 1124.232 | 255.210 | **<0.001** |
| Block 16-20 [1] | 0.979 | 0.001 | 0.976 – 0.982 | -14.016 | 256.985 | **<0.001** |
| Block 21-25 [2] | 1.007 | 0.001 | 1.005 – 1.009 | 6.003 | 289.047 | **<0.001** |
| Triplet Type [L H] | 0.997 | 0.001 | 0.996 – 0.999 | -3.501 | 274.785 | **0.001** |
| OCI-R score | 1.000 | 0.000 | 0.999 – 1.000 | -0.881 | 255.281 | 0.379 |
| Block 16-20 [1] x—Triplet Type [L H] | 1.002 | 0.001 | 1.000 – 1.004 | 2.493 | 110434.011 | **0.013** |
| Block 21-25 [2] x—Triplet Type [L H] | 0.998 | 0.001 | 0.997 – 1.000 | -2.063 | 110274.701 | **0.039** |
| Block 16-20 [1] x— OCI-R score | 1.000 | 0.000 | 1.000 – 1.000 | -1.676 | 258.715 | 0.095 |
| Block 21-25 [2] x— OCI-R score | 1.000 | 0.000 | 1.000 – 1.000 | 2.228 | 291.538 | **0.027** |
| Triplet Type [L H] x— OCI-R score | 1.000 | 0.000 | 1.000 – 1.000 | 1.345 | 277.633 | 0.180 |
| (Block 16-20 [1] x—Triplet Type [L H]) x— OCI-R score | 1.000 | 0.000 | 1.000 – 1.000 | -0.218 | 110426.496 | 0.827 |
| (Block 21-25 [2] x—Triplet Type [L H] x— OCI-R score | 1.000 | 0.000 | 1.000 – 1.000 | -1.284 | 110286.125 | 0.199 |
| **Random Effects** | | | | | | |
| σ^2^ | 0.030 | | | | | |
| τ_00_ _Participant_ | 0.007 | | | | | |
| τ_11_ _Participant Block 16-20 [1]_ | 0.000 | | | | | |
| τ_11_ _Participant Block 21-25 [2]_ | 0.000 | | | | | |
| τ_11_ _Participant Trplet Type [L H]_ | 0.000 | | | | | |
| ρ_01_ | 0.151 | | | | | |
|  | -0.008 | | | | | |
|  | 0.085 | | | | | |
| ICC | 0.196 | | | | | |
| N _Participant_ | 257 | | | | | |
| Observations | 110862 | | | | | |
| Marginal R^2^ / Conditional R^2^ | 0.004 / 0.200 | | | | | |

**Note.** The marginal R-squared considers only the variance of the fixed effects, while the conditional R-squared takes both the fixed and random effects into account. Degrees of freedom are based on Satterthwaite’s approximation. Statistically significant terms are highlighted in bold. Terms in brackets indicate the level of factor that is contrasted against the reference level

**Supplementary Table S9.** Results of the binomial mixed model on accuracy regarding ’New knowledge’ in Study 2. .

|  | **Prediction of accuracy** | | | | | |
| --- | --- | --- | --- | --- | --- | --- |
| *Terms* | *Log-Odds* | *SE* | *95% CI* | *z* | *df* | *p* |
| (Intercept) | 2.263 | 0.025 | 2.213 – 2.313 | 89.472 | Inf | **<0.001** |
| Block 16-20 [1] | 0.008 | 0.022 | -0.035 – 0.051 | 0.376 | Inf | 0.707 |
| Block 21-25 [2] | 0.100 | 0.018 | 0.064 – 0.135 | 5.457 | Inf | **<0.001** |
| Triplet Type [L H] | 0.035 | 0.013 | 0.009 – 0.061 | 2.664 | Inf | **0.008** |
| OCI-R score | -0.003 | 0.002 | -0.007 – 0.001 | -1.290 | Inf | 0.197 |
| Block 16-20 [1] x—Triplet Type [L H] | -0.005 | 0.017 | -0.038 – 0.029 | -0.271 | Inf | 0.787 |
| Block 21-25 [2] x—Triplet Type [L H] | 0.040 | 0.015 | 0.011 – 0.070 | 2.656 | Inf | **0.008** |
| Block 16-20 [1] x— OCI-R score | -0.001 | 0.002 | -0.005 – 0.002 | -0.736 | Inf | 0.461 |
| Block 21-25 [2] x— OCI-R score | 0.001 | 0.002 | -0.002 – 0.004 | 0.597 | Inf | 0.551 |
| Triplet Type [L H] x— OCI-R score | -0.001 | 0.001 | -0.003 – 0.001 | -1.141 | Inf | 0.254 |
| (Block 16-20 [1] x—Triplet Type [L H]) x— OCI-R score | 0.001 | 0.001 | -0.002 – 0.004 | 0.766 | Inf | 0.444 |
| (Block 21-25 [2] x—Triplet Type [L H] x— OCI-R score | 0.000 | 0.001 | -0.002 – 0.003 | 0.230 | Inf | 0.818 |
| **Random Effects** | | | | | | |
| σ^2^ | 3.290 | | | | | |
| τ_00_ _Participant_ | 0.126 | | | | | |
| τ_11_ _Participant Block 16-20 [1]_ | 0.041 | | | | | |
| τ_11_ _Participant Block 21-25 [2]_ | 0.021 | | | | | |
| τ_11_ _Participant Trplet Type [L H]_ | 0.008 | | | | | |
| ρ_01_ | 0.235 | | | | | |
|  | -0.161 | | | | | |
|  | -0.142 | | | | | |
| ICC | 0.043 | | | | | |
| N _Participant.Public.ID_ | 257 | | | | | |
| Observations | 122349 | | | | | |
| Marginal R^2^ / Conditional R^2^ | 0.004 / 0.047 | | | | | |

**Note.** The table shows regression coefficients of fixed effects and summary information about the random effects. Coefficients are log odds, thus positive values indicate that the given independent variable is associated with an increased likelihood of correct responses, and negative values are the opposite. P values for coefficients are from z tests, making the degrees of freedom infinite. The marginal R-squared considers only the variance of the fixed effects, while the conditional R-squared takes both the fixed and random effects into account. Statistically significant terms are highlighted in bold. Terms in brackets indicate the level of factor that is contrasted against the reference level.
